# Building from the ground up: augmentation strategies to improve protein language model performance on protein sets

**DOI:** 10.64898/2026.10.20.760553

**Authors:** Emily Stump, Hayden Dennison, Pietro Strobbia, George Stan, Ruxandra I. Dima

**Author notes:** co-first author (Emily Stump, Hayden Dennison).

## Abstract

An important underlying requirement for deep learning models is the availability of large datasets, however, in sciences such as chemistry and biology, generating experimental data of sufficient size represents a significant bottleneck. In this work, we probed the extent and nature of limited input data, characterize how several protein language models (pLMs) learned on suboptimal datasets, and investigated augmentation approaches to improve predictive model performance. We found that narrowing the distribution of protein-protein complexes in a limited dataset permits an order-of-magnitude decrease in training dataset size, if a model is only expected to generalize to in-distribution data. Similar results were found in active learning (AL), where both in- and out-of-distribution Matthews Correlation Coefficient (MCC) showed a small but consistent increase upon augmentation of the training set with complexes produced by ESM-3, a generative pLM.

## Introduction

As the size of modern neural networks increases to millions or billions of parameters, there is a corresponding increase to the need for data to train them.^1^ For domains such as natural language (particularly English), sufficient data can be obtained by leveraging the totality of written material available on the Internet, combined with label-free semi-supervised learning methods.^2^ In other domains, such as the chemical and biological sciences, there may not exist such a large corpus of data, presenting barriers to the successful adoption of deep learning in these fields.

The creation of labeled chemical datasets is generally a high-cost process, which requires both laboratory-grade materials and skilled human labor. ^3^ The recent boom in machine learning, revolving around data-hungry large language models (LLMs), puts greater focus on this fact.^4^ Although these models perform well in domains with high data availability, we have observed a lack of research performed on their capabilities and limitations in chemical research where data may be scarce.

Techniques such as active learning (AL) have been developed to address data scarcity by iteratively optimizing the creation of datasets under a limited budget.^5,6^ AL is a model-agnostic method that suggests how an existing dataset should be expanded to maximize the performance of downstream analyses, such as machine learning models, upon it. AL pipelines assume the existence of a so-called “oracle”, which can perform a labeling of any data point accurately but whose total number of labelings is limited. Thus, AL methods generally prioritize the labeling of data points that a downstream model predicts with low certainty, and de-prioritize labeling points that the model is already certain about (regardless of the accuracy of the predictions). Other variants exist that score prospective additions to the training set based on informativeness, redundancy with other prospects, etc.

AL has previously been explored in the context of chemical ML models, with a focus on small molecules and oligopeptides.^7,8^ Smith and Roitberg sought to generate datasets for the training of transferable machine learned interatomic potentials (MLIPs). ^9^ They selected molecules from external datasets that an ensemble of trained MLIPs disagreed upon when predicting the potential energy. By incorporating their additions, they were able to outperform a potential developed through random sampling with only a fraction of the training set size. An et al. used AL in the context of biomolecular condensates.^10^ They combined it with a generative procedure, using a genetic algorithm to create heteropolymers that exhibit phase separation for further study.

Other methods akin to AL have been applied successfully in other areas of computational chemistry. Molecular dynamics simulations often suffer from incomplete sampling of physically important regions of state space.^11^ Adaptive sampling seeks to address this limitation by performing an initial set of randomly initialized trajectories, then identifying the poorly sampled regions in order to construct seeds for additional trajectories. This iterative process can improve the exploration of state space, analogous to how AL can iteratively expand the training set to improve model performance. Li et al. demonstrated adaptive sampling in conjunction with ML methods to evaluate poor sampling, ranking the standard deviation of predictions from an ensemble of similar MLIPs to identify conformations for explicit quantum computation and subsequent addition to the training set.^12^

Other well-established techniques for supplementing insufficient datasets are over-sampling methods such as SMOTE^13^ and ADASYN,^14^ which are used to correct class imbalances by generating synthetic data samples. The goal of algorithms like these is to balance the ratio of minority and majority class samples, which can improve the downstream performance of ML algorithms. In an AL setting, over-sampling could be used to improve the predictive performance of any model that identifies uncertain data points, potentially making the calculation of uncertainty more reliable. However, synthetic over-sampling has limitations. Both ADASYN and SMOTE generate synthetic samples by interpolating between nearest-neighbors in the dataset, and ADASYN cannot be used with purely categorical data. Producing synthetic samples by interpolation is frequently appropriate for tabular data, but is difficult to apply when the data samples are variable in size, such as protein complexes.

Nevertheless, AL and over-sampling are no longer the only possibilities for dataset augmentation. There has been a rise in generative models that are trained on large datasets that scientists have been extending for years. For example, AlphaFold^15^ and ESM-3^16^ are protein language models (pLMs) trained on the recorded evolutionary record that can mimic the distributions of naturally occurring protein structures and sequences. Because it has been trained on such a vast corpus, ESM-3 is capable of producing biologically viable protein sequences and structures without requiring hands-on experimental labor. In certain problem domains these models are also able to provide training labels, allowing such models to serve as an oracle in and of themselves. In these contexts, AL becomes a method to engineer prompts for one model to generate new and helpful additions to another model’s training set.

LLMs can serve as *universal data augmenters* ^17^ that create human-like data for a variety of tasks, such as sentiment analysis and code generation. Data generated with LLMs has the advantage of being cost-efficient, scalable, controllable, and private. However, there are also challenges, such as ensuring synthetic data possesses sufficient variety without becoming out-of-distribution for a given task. One case study by Li et al. ^18^ found that a language model trained exclusively on raw data performed better across a range of tasks than a mix of raw data and synthetic data. However, the authors also observed a consistent performance increase by augmenting a small raw dataset with synthetic data, indicating that synthetic data can help address data scarcity in natural language processing tasks. The corpus of the English written language is mostly produced by human authors, while the language of proteins has been driven by stochastic evolutionary pressures.

In this paper, our focus was on evaluating whether uncertainty-based AL methods using pLMs could produce useful synthetic data for tasks in the chemistry domain. Specifically, we investigated best practices for protein data augmentation based on pLMs, the ability of the resulting augmented data to predict complex sequence-structure relationships in proteins, and the dependence of these findings on the extent and nature of limited input data. Because interactions between biomolecules lie at the core of most biochemical processes in cells, we selected as data source the DIPS-Plus dataset of protein-protein complex structures.^19^ This is a large dataset (*∼* 40,000 entries) of experimentally-determined protein-protein complexes and thus the number of elements in the set can be artificially reduced, or “ablated”, by removing points in a random, unbiased manner. Applying ablation allows us to explore how pLMs and AL behave on a range of sizes to mimic conditions of limited data availability.

To this end, we designed a neural net architecture that incorporated a pretrained pLM to predict the residue-residue contact map for a complex from the input of the sequences of the two protein chains. We explored pLMs ranging across parameter counts in the range of 10^6^-10^9^. The artificial reduction of the DIPS-Plus set mimics conditions of limited data availability. In realistic scenarios, however, one generally has to make a choice regarding the diversity of the examples in their dataset. If diversity is prioritized, the data points are thinly spread throughout the space of all possibilities; if it is not, the points will be clustered tightly into one region. This is expected to affect the learning capabilities of a model trained on the dataset, and so both size and scope are important factors to consider when building a dataset.^20^ Ideally one can create a dataset that is both large in size and wide in its scope, with the data points densely covering all regions of their space. An additional goal of this work was to explore the conditions where these desires cannot both be met.

The DIPS-Plus set was ablated to varying degrees, in a biased or unbiased manner, allowing us to determine how the pLMs tested performed on datasets differing in size and scope. The disagreement between the pLMs’ predictions was used to identify poorly understood complexes. We then used those complexes as seeds for the generative model ESM-3^16^ to produce additions to the training set.

Our work led to three key results: 1) sufficiently large pLMs can fail to achieve performance comparable to smaller pLMs if the training set is not large enough; 2) the performance of an appropriately sized pLM can be good even on a small dataset, so long as the problem scope is scaled to match the dataset size; and 3) the generative AL pipeline explored here performs best when the scope of the initial training set is small, so long as the model has proven capable of at least a modest level of learning on that initial set. These results provide important information on the capabilities of pLMs relative to data availability, as well as on when and how AL methods can be used to efficiently expand existing datasets. The generality of our findings likely makes them applicable beyond the realm of protein sets to wide varieties of LLMs and datasets.

## Materials and Methods

### Dataset

DIPS-Plus is a dataset comprising *>*40,000 protein-protein complexes.^19^ The target value of the dataset is the residue-residue contact map of each complex’s interface, where the contact is defined by the corresponding C*_α_*-C*_α_* distance within 6Å. Features annotated on the residue and atomic level are provided for each member, as well as their amino acid sequences. A training/validation (train/val) split is provided by the authors and was used here. In addition to the processing performed by the dataset’s creators, a check was performed on the DIPS-Plus dataset for any complexes repeated between the splits. These duplicates were removed to avoid cross-contamination between the training set and validation sets. For two items to count as duplicates, both of the sequences had to be duplicated as well as the contact map. Any duplicates were deleted from the training split, as it is the largest and so will be least affected by the reduction in size.

We assume that we can efficiently restrict the scope of the DIPS-Plus dataset by selecting an appropriate parameter and by only retaining complexes with a value for that parameter within some percentile range. In what follows we restricted our dataset using the buried surface area of the interface as said parameter. We also separately tested the use of the number of residue-residue pairs in contact at the interface as an alternative parameter to check for robustness. These parameters were chosen to serve as proxies for interface complexity. Interface area depends both on the number of contacts formed, and on the specific identity of the residues that form the interface. Larger residues contribute more area, and thus a larger and more diverse set of interactions that characterize the interface.

The selected parameter was used to ablate the training and validation sets to either 20%, 50%, or 80% of their original size, which we refer to as extensive, moderate, or limited ablations. Each of the three levels of ablation was performed in two distinct ways. In the uniform style of ablation, the values of the chosen parameter (interface area or residueresidue pair count) across the dataset were separated into 10 bins and stratified sampling was applied. The end result was a randomly sampled subset that maintains the distribution of the parameter. We considered this type of ablated set to be broad in the DIPS-Plus space but sparse in its coverage. In the so-called targeted (or biased) style of ablation, whichever parameter was used was restricted to a randomly placed 20, 50, or 80 percentile range. The percentile ranges were calculated using the complete dataset (train and validation splits together). For a particular targeted ablation, both training and validation sets are restricted to the same percentile range. These ablated sets are viewed as being narrow in their spread but densely covered.

For each subset size (20%, 50%, or 80% the full dataset) and ablation style (uniform or targeted), 5 independent ablations were performed for a total of 30 different subsets for residue-residue pair count and 30 different subsets for interface area used as the ablation condition variable. In addition, we held out a small, randomly selected portion of the validation set to use as an unseen test set characteristic of the complete DIPS-Plus dataset distribution. This held-out test set was the same for every trial and was not subject to any form of ablation. We use the term “validation set” to refer to what remains of the validation set after the test/hold-out set was sampled and set aside.

Processing the dataset and performing ablations was done in Python using a combination of the Pandas, NumPy, SciPy, and scikit-learn libraries, and the models were trained in PyTorch.^21–25^

### Models

We ignored the residue- and atomic-level features available in DIPS-Plus, only using the sequences of the chains in each complex to represent each element of the dataset. The goal of each model we constructed was to predict a complex’s residue-residue contact map given only these two sequences. The model architecture began with a twin pair of pretrained BERT-style encoders^2^ that took one of the two sequences as input and outputted a sequence of D-dimensional residue-level contextual embeddings for that sequence 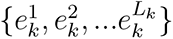, where *k* denotes either sequence 1 or 2 of the pair, and *L_k_*is the total length of the sequence. These two sets of residue embeddings were assembled into an array *M* defined as *M_ij_*= *concat* 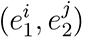, where *concat* is the operation of concatenating the second argument with the first to form a single vector of length 2 *∗ D*. This results in a *L*_1_ *× L*_2_ *×* (2 *∗ D*) array, with the (*i, j*)th element representing the pair of residue *i* from the first sequence and *j* from the second.^26^ A multilayer perceptron (MLP) with a single sigmoid output was applied positionwise across the 2 *∗ D* length set of residue embedding pairs. The output of this MLP is the probability of those two input residues forming a contact. Using a threshold of 0.5 to take this probability to 0 or 1, the model outputs an *L*_1_ *× L*_2_ size residue-residue contact map for the protein-protein pair.

Each residue-residue pair was treated as a single prediction, allowing us to use metrics for binary classification in a position-wise manner and then average them across the whole *L*_1_ *× L*_2_ map of predictions to achieve a single value for the input complex. Binary cross entropy (BCE) was used in this manner as the loss for training this network, while Matthews correlation coefficient (MCC) was similarly used to evaluate the performance of trained models.

Three different BERT-style models of increasing size were tested as the encoder pairs.^27–29^ This allowed us to form a set of models that could later be used for our AL protocol, further discussed below. These models are summarized in Table 1, and further details regarding pretraining, architecture, etc. may be found in their associated papers. Note that the authors of ESM-2 made available several versions of the model, varying in size from 8 million to 15 billion trained parameters. We chose the 8M parameter model for use in this work, and all further references to ESM-2 should be understood as referring to that specific model. The model weights for ESM-2 and ProtBERT were downloaded from Hugging Face,^30^ while AminoBERT’s came from its GitHub page.

**Table 1:** pLMs used in this study, along with their approximate parameter count and architectural details.

| Model | # Parameters | # Attention heads | Embedding dimension |
| --- | --- | --- | --- |
| ESM-2 (8M) | 8M | 20 | 320 |
| AminoBERT | 87M | 12 | 768 |
| ProtBERT | 420M | 16 | 1024 |

Hyperparameter optimization (HPO) for each model was performed using the Optuna package for Python.^31^ The hyperparameters tested, as well as the values explored are listed in Table S1. The architectural hyperparameters (depth, width, activation function, and dropout rate) pertain only to the positionwise MLP. A position weight was used for the BCE loss because contact maps are fairly sparse, creating imbalance in the classes when viewing each position in the *L*_1_ *× L*_2_ array as a separate binary prediction.

For each HPO trial, the model was trained on the complete train set, and the set of parameters with the best MCC on the validation set was used for training that model on every ablation trial. This was done to avoid the computational requirements of performing HPO for every single ablated dataset.

### Measuring the similarity of contact maps

Due to the nature of the results detailed below, we used only AminoBERT and ESM-2 for our AL protocol. In order to measure the agreement between the protein-protein contact maps predicted by the two models, we utilized the central kernel alignment (CKA) method. The contact maps predicted by an AminoBERT and an ESM-2 model for a protein complex *i* are denoted by **A***_i_* and **E***_i_*, respectively. The (linear) CKA-score for two given predictions of the same protein complex *i* is then given by Equation 1 where **K***_A_* denotes the linear kernel for an AminoBERT model’s prediction and **K***_E_* the linear kernel of an ESM-2 model’s prediction.

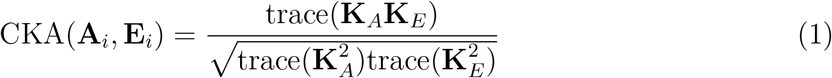

The general form of the linear kernels is 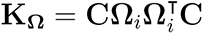. In this equation, **C** is a matrix that performs a mean-centering operation by column, defined as 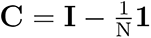, with N being the number of rows in the matrix **Ω***_i_* = *{***A***_i_,* **E***_i_}*. **1** is an N *×* N matrix where every element is the value 1, here the same number of rows and columns as **C**. We note that the contact maps predicted by AminoBERT and ESM-2 are continuous values between 0 and 1, with each element being the probability of a contact forming.

The CKA-score for two contact maps is a number between 0 and 1, with 0 representing no alignment between the centered kernels and 1 representing perfect alignment (i.e., the two contact maps are identical). A protein complex having a CKA-score close to 1 indicates that the two models AminoBERT and ESM-2 produced similar contact maps for that protein complex and can be said to “agree” on it. Conversely, complexes with CKA-scores close to 0 indicate that the two models disagree on their predictions for that protein complex.

We evaluated three other measures of contact map similarity: the Singular Angle Similarity,^32^ the Frobenius matrix norm of the difference between two contact maps, and the nuclear norm of the difference between two contact maps.^33^ Representative results are shown in Figure S1. The distributions of CKA-scores exhibited the least overlap between the train, validation, and test sets, which informed our choice to use the CKA-scores for distinguishing between protein contact map representations.

### Selecting disagreed-upon protein complexes and perturbing their sequences

Once the uncertainty of each protein complex under a given ablation is computed, we select the complexes from the 20th percentile of validation set CKA-scores scores as the basis for generating new protein complexes for each trial. The number of complexes found to be disagreed-upon are shown for each ablation in Table 2.

**Table 2:** The number of protein complexes in the 20th percentile of validation set CKA-scores for each ablation. Uncertainty reported is standard error of the mean.

| **Ablation condition** | **Ablation variable** | **Ablation degree** | **Uncertain complexes** |
| --- | --- | --- | --- |
| uniform | residue-residue count | extensive | 1267 |
| uniform | residue-residue count | moderate | 3163 |
| uniform | residue-residue count | limited | 5059 |
| targeted | residue-residue count | extensive | 1320 ± 20 |
| targeted | residue-residue count | moderate | 3261 ± 22 |
| targeted | residue-residue count | limited | 5126 ± 6 |
| uniform | interface area | extensive | 1267 |
| uniform | interface area | moderate | 3163 |
| uniform | interface area | limited | 5059 |
| targeted | interface area | extensive | 1284 ± 12 |
| targeted | interface area | moderate | 3192 ± 15 |
| targeted | interface area | limited | 5055 ± 8 |

We masked approximately 20% of each protein complex sequence with the ESM-3 mask token (“ ”) and prompted ESM-3 with the masked sequence to generate a novel protein complex.

### Masking dissimilar protein complexes

Table 3 shows the probabilities used to scramble the amino-acid sequences of the dissimilar protein complexes. For each amino acid in the seed complex, the masking algorithm randomly selects an operation between 0 and 4 inclusive according to probabilities conditioned on whether the prior operation was or was not either of the masking operations (operation ID 0 or 1). This was done to encourage continuous stretches of masked amino acids. The measured percent of each protein sequence masked is shown in Figure S2.

**Table 3:** Probabilities of each operation when iteratively masking protein complexes to prompt ESM-3. Only operations 0 and 1 cause the probabilities to be selected from the “prior operation masking” column. Residues were mutated by randomly selecting a residue according to their relative abundance in the human proteome.^34^

| operation ID | description | prior operation<br>not masking | prior operation<br>masking |
| --- | --- | --- | --- |
| 0 | mask position | 0.100 | 0.30 |
| 1 | insert extra mask<br>token | 0.010 | 0.20 |
| 2 | remove current<br>token | 0.010 | 0.20 |
| 3 | mutate residue | 0.001 | 0.00 |
| 4 | perform no<br>operation | 0.879 | 0.30 |

### Generating novel protein complexes by unmasking

The scrambled dissimilar protein complexes found for each ablation scheme were used to generate novel protein complexes with the ESM-3 generative protein language model. ^16^ We used ESM-3’s guided generation feature to drive generated protein complexes towards high predicted protein template modeling (pTM) scores to ensure they were of sufficient quality to be used in the training process. Briefly, the pTM score compares a structure to a library of known quality structures.^35^ It is defined in equation 2:

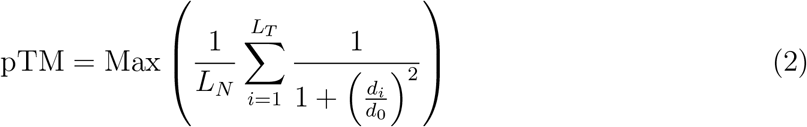

where *L_N_*denotes the number of residues in a generated or native protein structure, *L_T_* is the number of aligned residues to a template structure, *d_i_* is the distance in Å between two aligned residues, and *d*_0_ is a normalizing constant in Å equal to 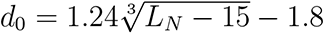 pTM scores range between 0 and 1 inclusive. A pTM score below 0.2 indicates that the protein structure has a low match to any template structure (0.17 pTM is the average for any two random protein structures), implying the generated protein complex possesses structural defects or is unrelated to the seed protein complex. The distributions of pTM scores for the generated proteins from interface area ablation are shown in Figure 1. We note that a small number of the generated proteins have pTM scores below 0.17, but the vast majority have pTM scores above 0.2. Compared to the distributions of pTM scores for the DIPS-Plus dataset (Figure 2A), these scores are generally smaller and with less variance. The ESM-3 model can use multiple input tracks to constrain the physical properties of its outputs. We used a generation temperature of 0.3 and unmasked 3 tokens for every inference step. The default generation temperature is 1.0 and the default number of tokens to unmask per step is the number of total masked tokens divided by 20. We instead chose to use an iterative generation strategy (which often results in higher-quality final structures) to unmask 3 tokens per step for every protein regardless of length. The generation temperature controls the “creativity” of the generation process by causing the model to fill in each mask with alternative tokens than the highest probability token. The secondary structure annotations and solvent-accessible surface area values provided in DIPS-Plus were used as additional input tracks, masking the same positions as the amino acid tokens, in a manner similar to the out-of-distribution guided inference experiment by the ESM-3 authors. Any values missing in the dataset were replaced with <pad> tokens. Two examples of protein complexes that had low similarity scores for the interface area 20% data retained targeted ablation are shown in Figure 3, along with the generated protein complexes that arose from unmasking their sequences. One selected protein, with Protein Data Bank (PDB) ID 3ODH, is a restriction enzyme and the other, with PDB ID 1ORU, is an uncharacterized protein.

**Figure 1:**
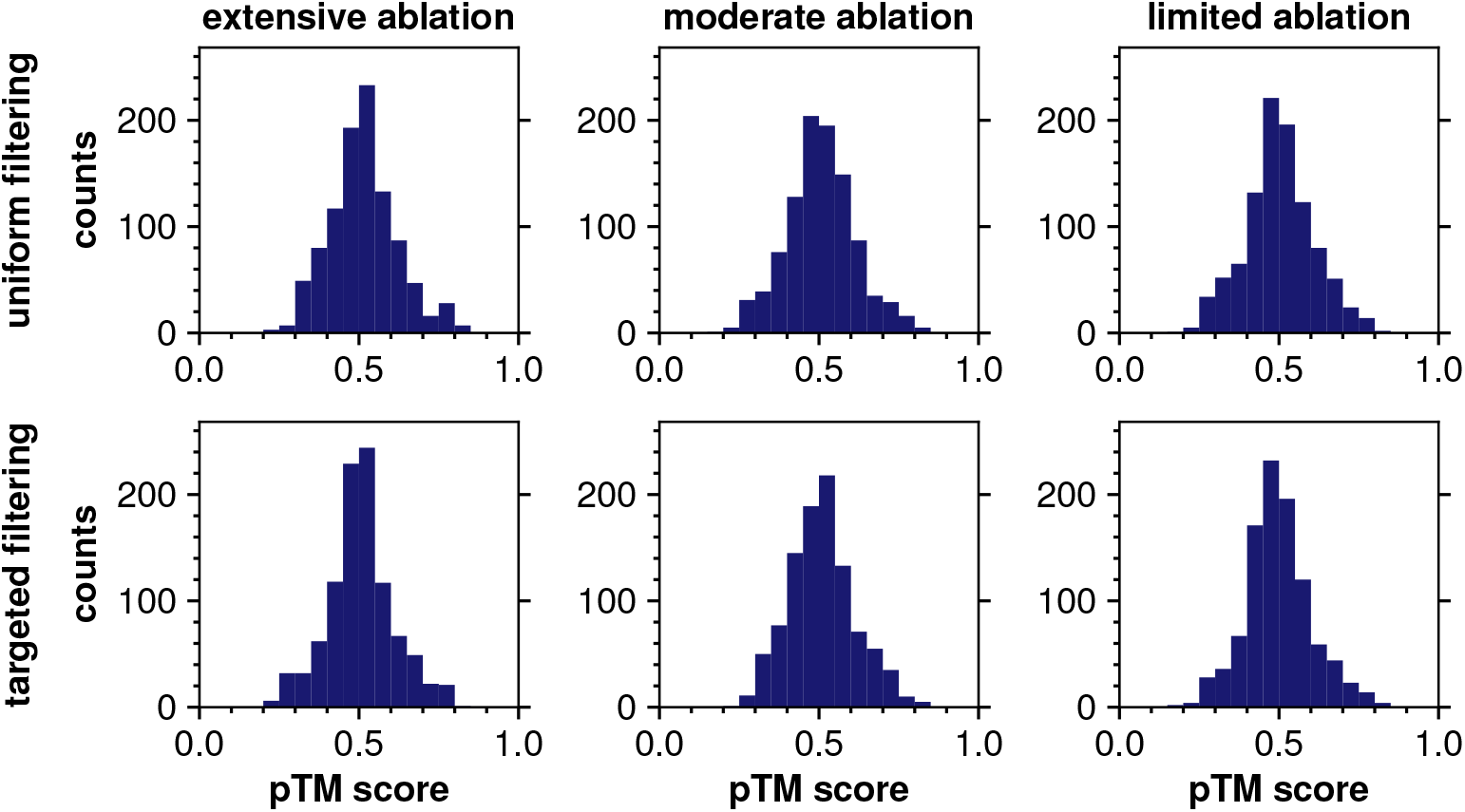
Quality of ESM-3 generated proteins using the interface area as ablation variable. Distribution plots of the pTM scores are shown for uniform (top) and targeted (bottom) filtering performed using extensive (left), moderate (middle), and limited (right) ablation. All distributions have a mean value near 0.50 pTM and a standard deviation of ~ 0.01. The mean pTM score for our generated proteins is similar to the mean value of 0.52 ± 0.10 for ESM-3 with unbiased prompting.^16^ Bin width is 0.05.

**Figure 2:**
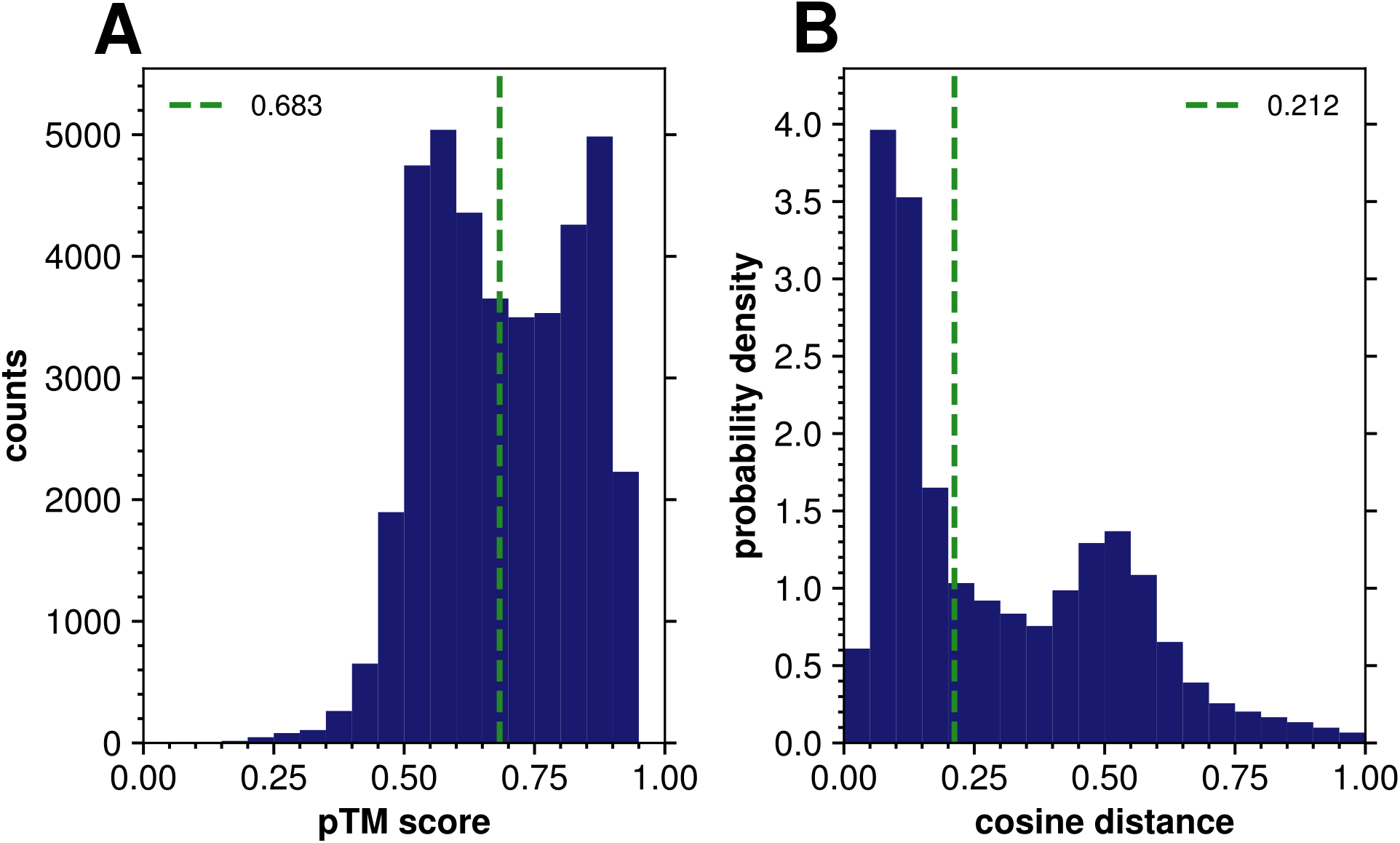
Quality of the DIPS-Plus protein complexes according to pTM score. Histograms of the distribution of (A) pTM scores and (B) cosine distances between embeddings of the DIPS-Plus proteins are shown. In both panels the median value is indicated with a vertical green line. Bin width for both panels is 0.05.

**Figure 3:**
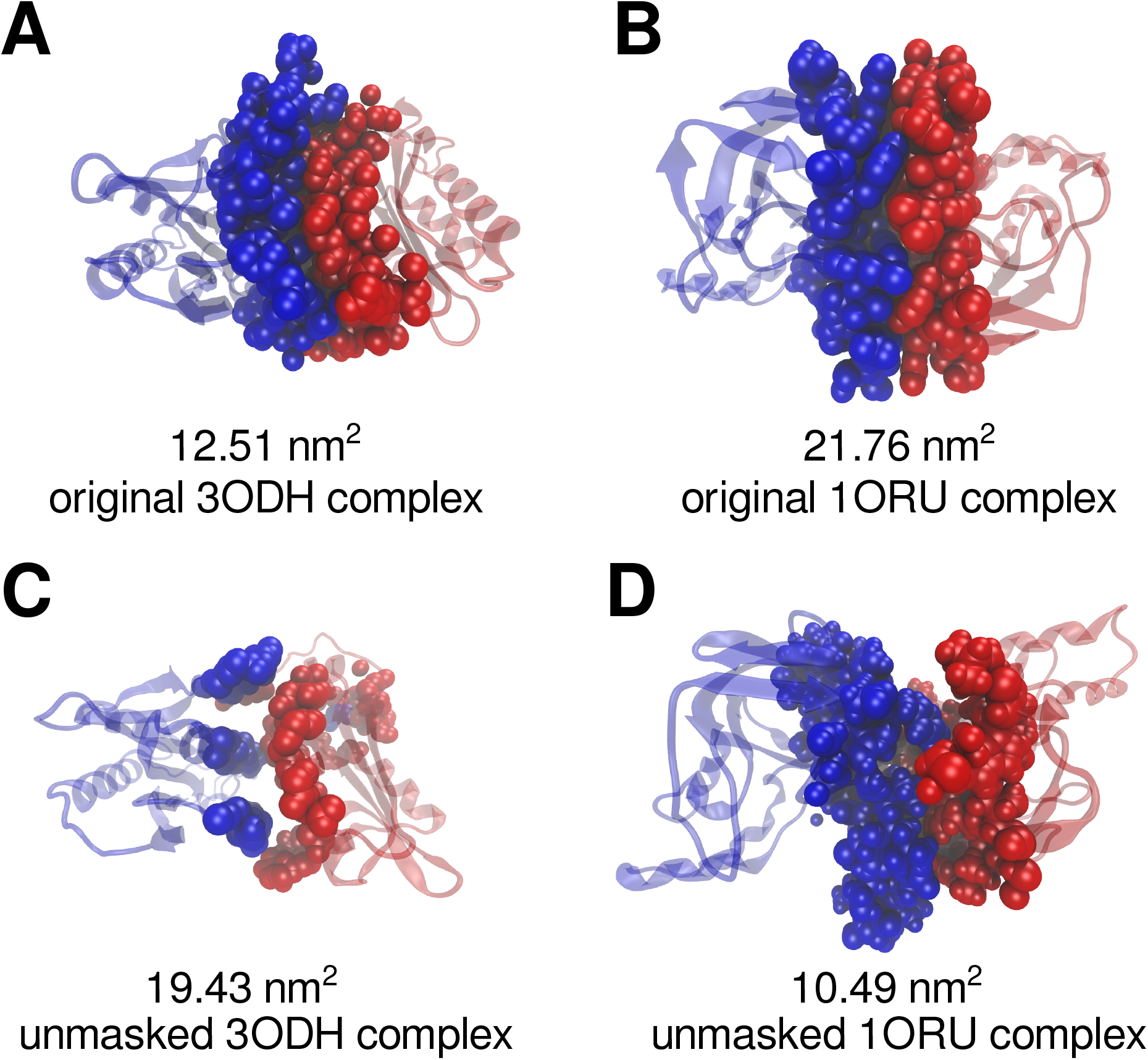
Examples of structural differences between generated complexes and their seeds. (A) (B) The original (seed) three-dimensional structures of complexes with Protein Data Bank IDs (A) 3ODH and (B) 1ORU, respectively. Both complexes have low similarity contact map predictions (CKA-scores scores of 0.084 for 3ODH and 0.138 for 1ORU). (C) (D) The generated protein complexes for (C) 3ODH, and (D) 1ORU. For all structures, chain A is colored red and chain B blue. Residues within 0.9 nm^2^ of the interface are shown by using the van der Waals representation. The protein complex generated from 3ODH (C) has a qualitatively different interface from its seed, while the protein complex generated from 1ORU (D) has a reduced but similar interface.

### Measuring distances between protein complexes in latent space

We used the ESM-model^27,36^ with 6 billion parameters to embed all generated protein complexes in the same latent space as the DIPS-Plus complexes. In this latent space, the distance between two protein complexes is the cosine distance between their embedding vectors *v*_1_ and *v*_2_, defined as

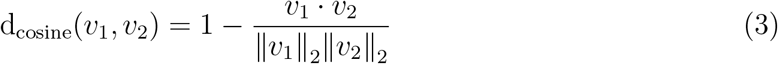

The distribution of the cosine distances for the entire DIPS-Plus (Figure 2B) is characterized by a relatively small median value of 0.212, indicative of the fact that a sizable group of the protein complexes clusters together in embedding space.

## Results and Discussion

### Effects of Dataset Ablation on Model Performance

We started by evaluating the ability of the various pLMs to predict the contact maps of protein complexes based either on the full DIPS-Plus dataset or the subsets obtained by different ablation approaches. For each combination of model (ESM-2, AminoBERT, ProtBERT), ablation condition (either uniform or targeted ablation to 20, 50, or 80% of the full dataset’s size), and ablation variable (residue-residue pair count or interface area), the train, validation, and test MCCs were averaged across the five trials. These mean MCCs for the ablations based on interface area, along with standard error bars and the MCCs on the full dataset are presented in Figure 4. The results for residue-residue pair count-based ablations are shown in Figure S3, and are similar to the ones discussed here.

**Figure 4:**
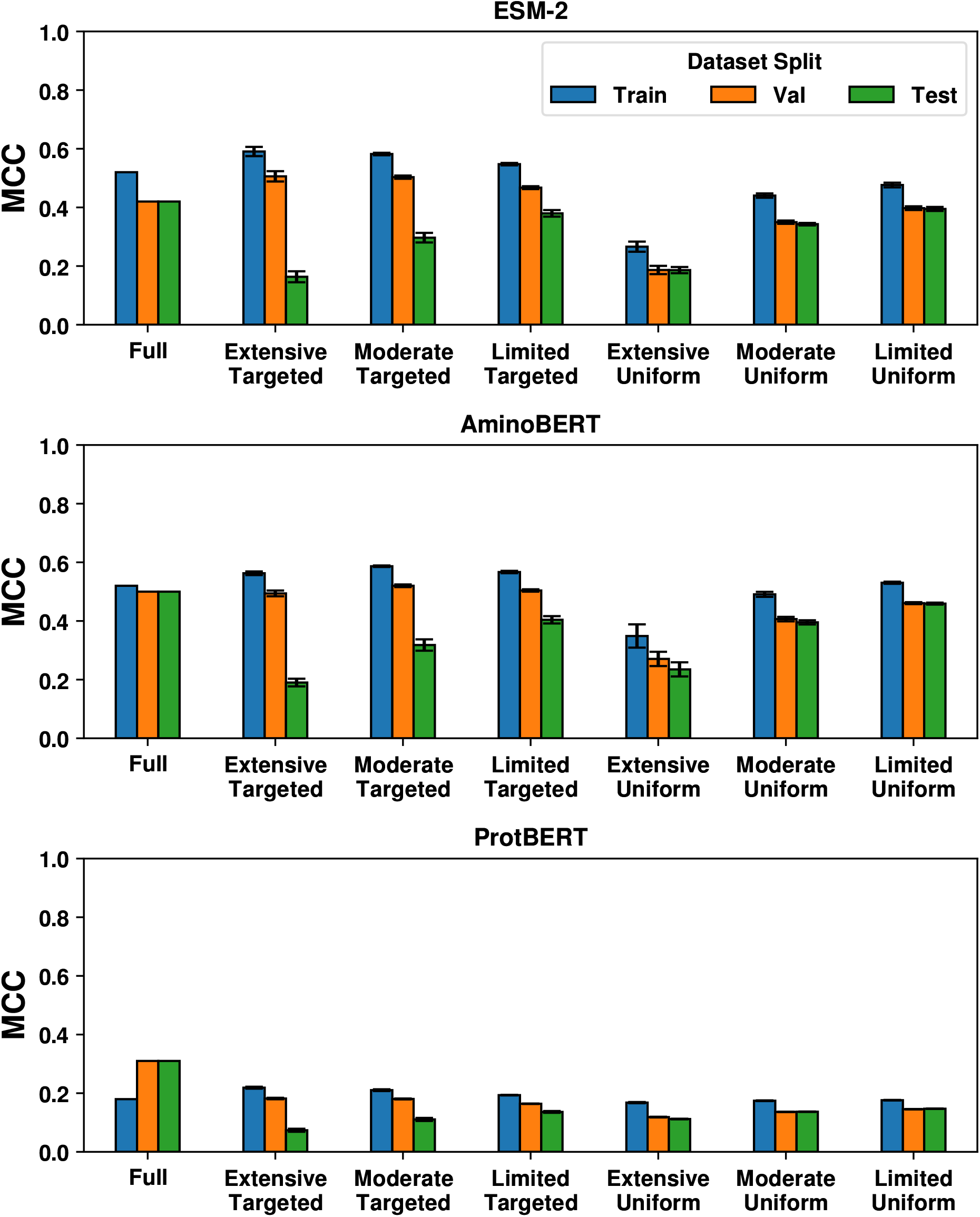
Performance of pLMs on DIPS-Plus data. Bar plots indicate the average MCC values and standard error for each pLM tested (ESM-2, top; AminoBERT, middle; ProtBERT, bottom), across the five trials performed, for the train (blue), validation (orange), and test (green) sets for a given ablation condition. Corresponding MCCs for the full DIPS-Plus dataset are also provided for each pLM.

First, we make note of the disparity in performance between ProtBERT and the other two models. Average MCCs of ProtBERT models struggle to surpass 0.2 even on the training set, while the MCCs of the other models can get as high as 0.5. Note that ProtBERT was the largest model tested, with 420 million parameters. This may imply that the DIPS-Plus dataset is too small for a model of that size to effectively train on.

We found that training and validation MCC both benefit from a focused distribution compared to a broader one. For targeted ablations, these scores are consistently higher than those for uniform ablations of the same size. We also see negligible dependence of train and validation MCCs on the size of the dataset in the focused ablations. This is in contrast to the uniform ablations, where train and validation MCCs increase in concert with the size of the training set. In fact, an 80% retained uniform ablation leads to lower train and validation MCCs than the much smaller train set of the 20% retained targeted ablation.

The test MCCs, on the other hand, are consistently worse for targeted ablations compared to uniform ones. This demonstrates the importance of data distribution compared to dataset size, as the uniform ablations were meant only to decrease the size of the training set without affecting its distribution. This is also why the validation MCCs are nearly equal to the test MCCs of the uniformly ablated trials. As the size of the ablated sets increases, the test MCC increases in concert. The difference between the test MCCs for uniform and targeted ablations are not as extreme as one might expect, however, with the difference being within 0.1. Comparatively, the amount by which train and validation MCCs for targeted ablations are greater than those of uniform ablations is as high as 0.32 in some cases.

We also note how the MCCs of the targeted and uniform ablations relative to those on the full dataset change with respect to the degree of ablation. Targeted ablation to 20% of the full dataset shows the best train and validation MCCs, even exceeding those found on the complete dataset. As the degree of ablation decreases, these MCCs decrease to be closer to the full-dataset values. The test MCCs, conversely, increase in value. Uniform ablation to 20% of the full size, on the other hand, has worse train, validation, and test MCCs than those on the full dataset. As the degree of ablation becomes less severe, these MCCs increase steadily, with uniform ablation to 80% showing comparable performance to when the model has the full dataset available. Uniform ablation did not change the distribution of the data shown to the model, and these results demonstrate that additional data helps the model to develop a greater understanding of it. The results for targeted ablation demonstrate that the reduced dataset size can be compensated for by reducing the scope of the problem. We do, however, make note of the margin between validation and test MCCs being much greater for the targeted ablations than for the full set or the uniform ablations. While the model is more than capable of learning the restricted distribution despite a smaller amount of data, especially in the extensive ablation case, it severely overfits to it and is unable to generalize to the full distribution represented by the unablated test set. Additional insight into the differential behavior of models trained on targeted versus uniformly ablated sets are provided next.

### Models trained on datasets obtained with targeted ablation agree more strongly than models trained under uniform ablation

We measured the performance of the AminoBERT and ESM-2 models from each trial on the held-out test set. We found that the test set MCCs for models trained under uniform ablation are generally larger than those for models trained under targeted ablation of the same degree, whereas the opposite is true for the train and validation sets MCC. However, the AL protocol described in the Methods is based on model disagreement as measured by CKA, not predictive accuracy. Plotting the CKA-scores (Eq. 1) for the predicted complexes in each set shows similar trends to the MCC scores, with ESM-2 and AminoBERT predicting more similar contact maps on the train and validation set under targeted than uniform ablation. The as evidenced by the peak of the respective CKA distribution located at values closer to 1 (Figure 5). Conversely, the models agree more on the test set under uniform than targeted ablation. We also note that the distributions for all three sets converge to a similar shape as the degree of ablation decreases from extensive to limited.

**Figure 5:**
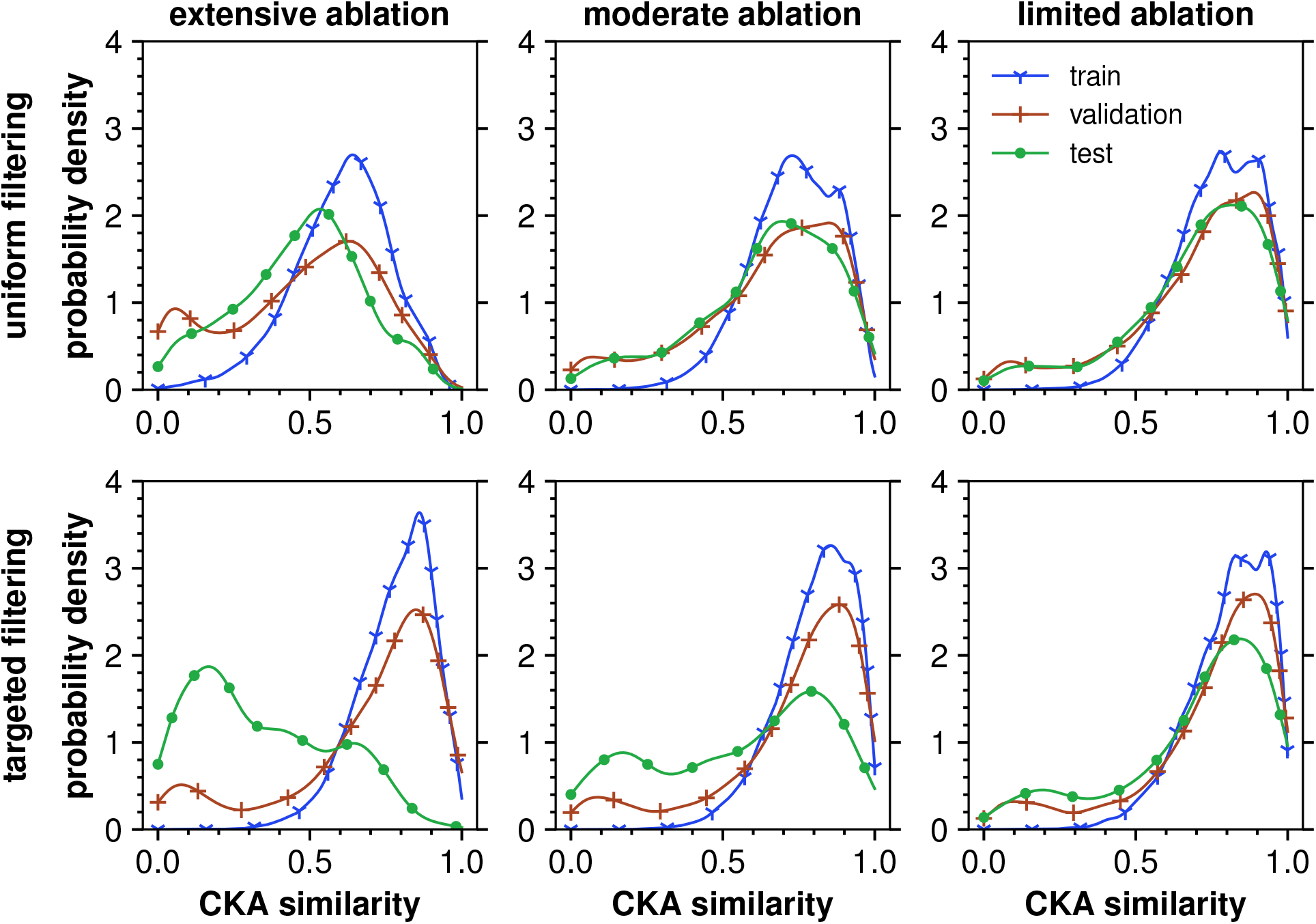
CKA similarity of ESM-2 and AminoBERT models on ablated data. Gaussian kernel density plots of CKA-scores are shown for train (blue), validation (red) and test (green) sets averaged across all 5 trials for each of uniform (top row) and targeted (bottom row) ablations of extensive (left), moderate (middle) and limited (right) degree, where ablation was done using interface area.

The train set CKA distributions under targeted ablation are more negatively skewed than the corresponding uniform ablation distribution at the same ablation degree (see Figure 5 and S4). However, for the test set, the uniform ablation distributions always exhibit negative skew, while the test set CKA-scores after extensive targeted limited ablation exhibit positive skew (Figures S5 and S6). We infer that, as the degree of ablation increases, targeted ablation results in a train set that diverges dramatically from the test set while exhibiting better MCC and CKA-scores. Reducing the number of training examples in a targeted way shrinks the variance of the distribution a model has to learn and allows it to perform better in that small protein space due to a higher bias.

### Characterizing similar and dissimilar complexes

Next, we focused on characterizing the differences between protein complexes for which the pLMs agree, versus the ones where they disagree in the predictions. Specifically, we characterized complexes from the validation set with high and low disagreement between the models’ predictions under different ablation conditions. The DIPS-Plus set itself is highly skewed with regards to what interface areas are represented within it, with strong peaks at *∼* 5.5 and *∼* 13 nm^2^, and the density dropping off beyond *∼* 20 nm^2^ (see the black kernel density plot in Figure 6). Model disagreement in ablations that cover most or all of this complete range is strongly connected to this inherent modality of DIPS-Plus. Trial 3 of the 80% targeted ablation demonstrates this, with the similar complexes of the validation set (in the top 20% of CKA scores) clustering around the existing modes, and dissimilar complexes (in the bottom 20% of CKA scores) clustering between and beyond these modes (see the orange vs. the blue density plots in Figure 6). Importantly, this behavior is seen in all of the uniform ablations (see supplementary material). Figure 6 shows examples of a similar and dissimilar complex selected from this trial. The similar complex, PDB:1MN7, has a calculated interface area of 6.28 nm^2^ and a CKA score of 0.93. The dissimilar complex shown, PDB:5A9D, has an interface area of 16.88 nm^2^ and a CKA score of 0.003.

**Figure 6:**
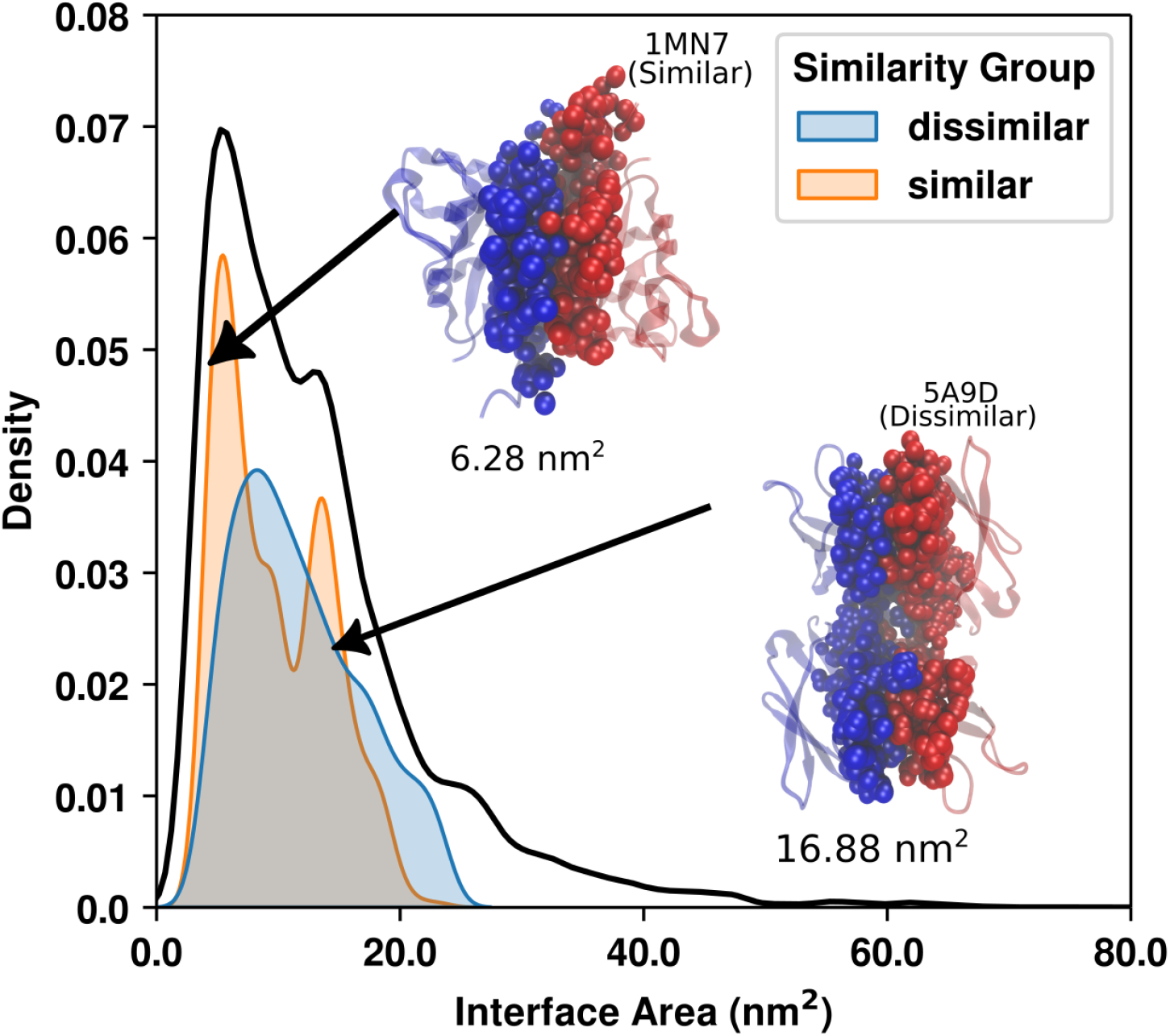
Characterization of similar and dissimilar complexes in limited, targeted ablation. Interface area distribution for the similar (orange, filled) and dissimilar (blue, filled) complexes identified in the third trial of limited, targeted ablation. The interface area distribution for the complete DIPS-Plus set (black, unfilled) is shown as well. Approximate locations of representative complexes described in main text are indicated with arrows.

The second trial of the 20% targeted ablation, in contrast, only contains complexes with interface areas between *∼* 7 and 13 nm^2^, i.e. between the two modes of the full DIPS-plus set (see Figure 7). In this situation where the range is more uniformly covered, we found clear distinctions between the two classes. The distributions of the interface area show that dissimilar complexes have a tendency towards the extreme ends of the sampled range, while similar complexes favor the middle values (compare the blue with the orange density plots in Figure 7). An example of a dissimilar complex for this 20% targeted trial is found between chains A and B in PDB:1Y13. This had a calculated interface area of 7.92 nm^2^, and the CKA between the contact maps predicted by AminoBERT and ESM-2 was 0.02. We also show an example of a dissimilar complex from the other end of the sampled interface area range, that being between chains B and C of PDB:3QID, with interface area 11.17 nm^2^ and a CKA of 0.0005. An example of a similar complex in this trial was between chains B and D of PDB:4KWG, with an interface area and CKA of 9.78 nm^2^ and 0.97, respectively. These complexes are shown in Figure 7, with the interfacing residues depicted as opaque van der Waals spheres and the remainder as a translucent cartoon.

**Figure 7:**
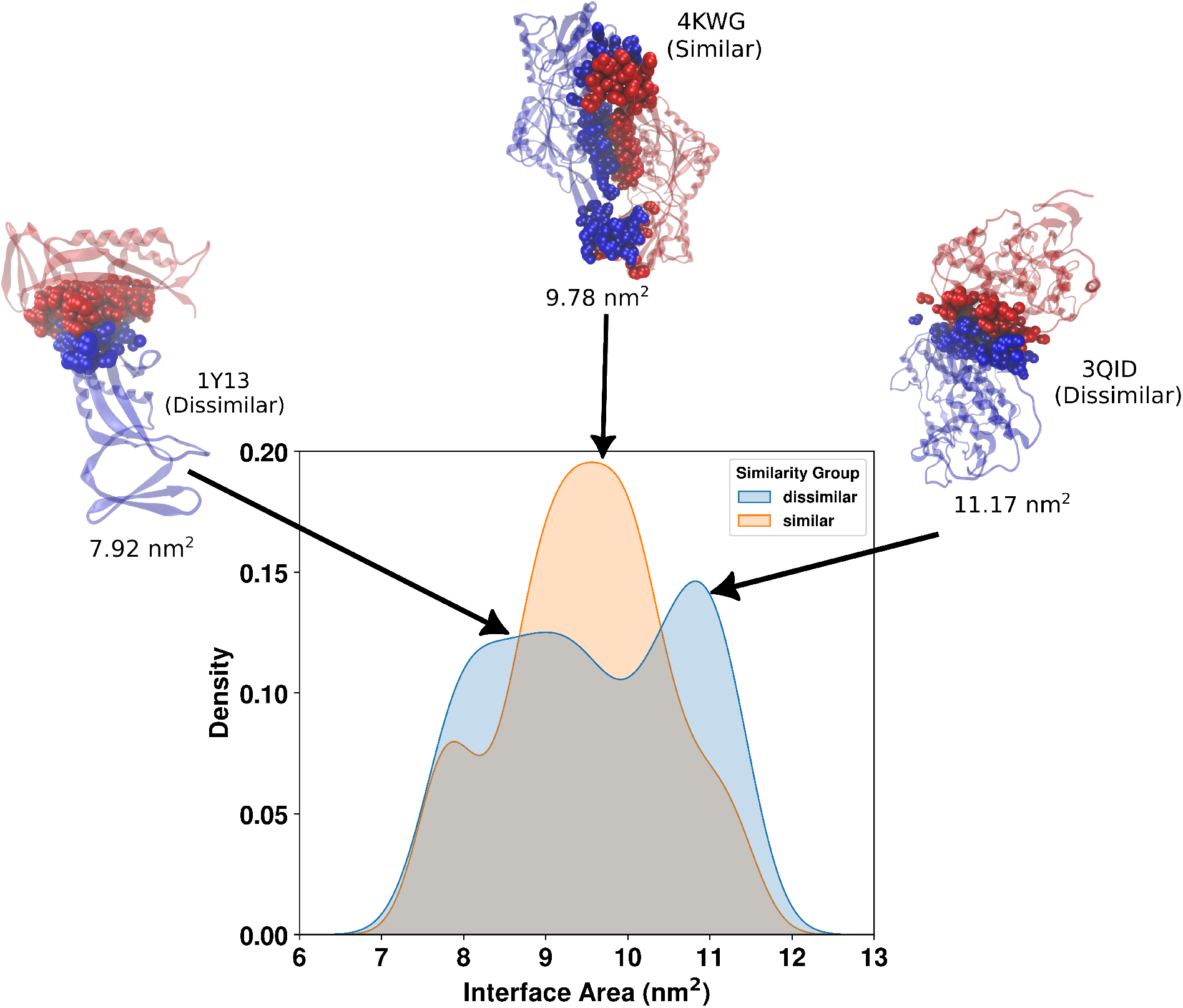
Characterization of similar and dissimilar complexes in extensive, targeted ablation. Interface area distributions for the similar (orange) and dissimilar (blue) complexes identified in the second trial of extensive, targeted ablation. Approximate locations of complexes described in main text are marked with arrows.

The interface area distributions of similar and dissimilar complexes for all trials are shown in Figures S7 and S8. Those figures display the distributions in terms of count, not density. This is to highlight both the spreads of each trial’s ablated set and their relative sizes.

The above results demonstrate both the robustness of using interface area as a way to group the DIPS-Plus set, and what factors affect model disagreements. Model disagreement appears to be driven most strongly by the underlying distribution, with models agreeing more readily on parts of data-space that have been well sampled. The modes of the full DIPS-Plus set tend to be where similar complexes are located, while dissimilar complexes are found between these modes and in the long tail of the distribution. When there is no clear enrichment in one region or another due to the placement of the percentile range sampled, dissimilar complexes instead cluster to the ends of the sampled range.

#### Ablated protein complexes are more likely to be near disagreed-upon than similar complexes

Beyond the insight into the interface area propensity of the similar versus the dissimilar complexes provided above, we were interested in characterizing the degree to which these two sets reside near the removed protein complexes in latent space. To use dissimilar complexes to augment an ablated dataset, we expect that they, rather than the similar complexes, reside closer to the removed proteins that we are trying to compensate for in driving model learning. To this end, we plotted the distributions of distances between the embeddings of protein complexes for which the two models AminoBERT and ESM-2 disagreed and those protein complexes which were removed as part of the ablation process. The dissimilar complexes are from the bottom quintile of CKA-scores similarity scores on the validation set contact maps for each ablation. The similar complexes are from the top quintile of CKA-scores. Figure 8 shows that, regardless of the percentage of data retained, the protein complexes for which the two models AminoBERT and ESM-2 disagree are closer on average to removed complexes in ESM-C embedding space than the complexes for which the models agree. These distributions for the ablations based on residue-residue count are shown in Figure S10.

**Figure 8:**
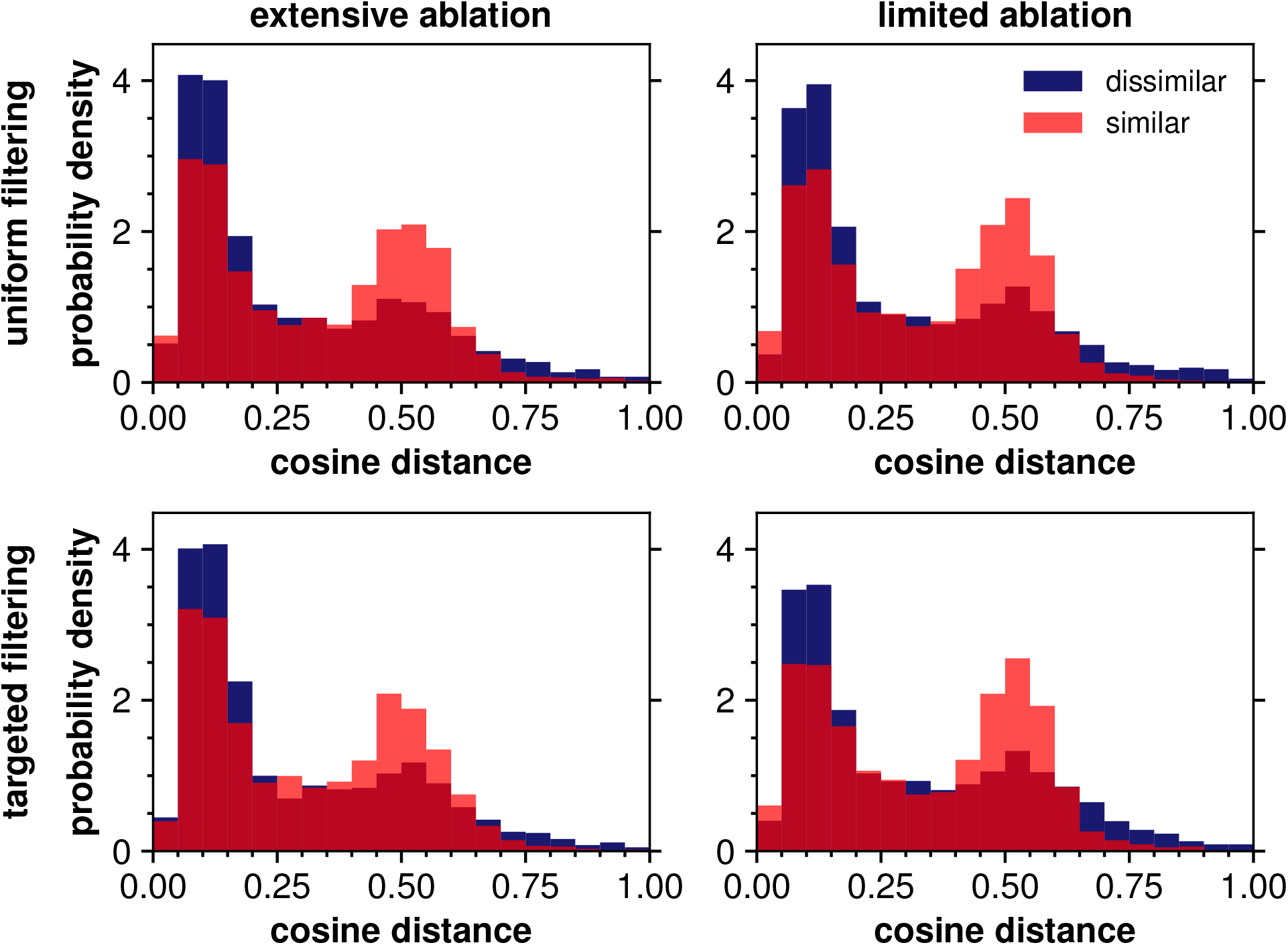
Embedding-space proximity of similar and dissimilar complexes to those removed in ablation. Distributions of cosine distances between dissimilar (blue) and similar (red) protein complexes vs removed proteins under uniform (top) and targeted (bottom) interface area ablation of extensive (left) and limited (right) degree. The dissimilar complexes are closer overall to the ablated complexes than the similar complexes. The embeddings were obtained from the ESM-C 6B model.^36^ Binwidth is 0.05.

In comparison to the overall DIPS-Plus cosine distribution (Figure 2B), both the dissimilar and similar embedding distributions are bimodal with the same peaks around 0.1 and 0.5 cosine distance. However, the similar distributions have a larger median value on average (0.320) than the entire DIPS-Plus set and the dissimilar distribution has a smaller median value (0.195), indicating that the top quintile and bottom quintile of CKA-scores can be used to find areas where data is missing from the datasets given to the models. We note that the Jensen-Shannon (JS) distance between dissimilar and similar distributions is smaller on average for the targeted ablations than the uniform ablations, which indicates that AminoBERT and ESM-2 models trained after targeted ablation can better identify out-of-distribution samples in their training sets.

### Distributions of generated protein complexes are biased toward the training distribution of the generative model

Ideally, the distribution of training data for a machine-learning model matches the distribution of the true population that produced the data. We examined how perturbing and ablating the distribution of training data for a set of models influences the confidence in that set’s predictions. But can a set of models trained on a distribution of data significantly different from the true population be utilized to identify the missing data points? We observe that the cosine distances between the embeddings under uniform and targeted ablation exhibit the same profiles as the training dataset is ablated (Figure 9). We infer that because ESM-3 has been trained on all currently available protein structures and sequences, of which DIPS-Plus is a subset, proteins it generates are biased toward the center of ESM-3’s training distribution, not the entire DIPS-Plus distribution, nor the ablated training distributions. In Figure 10, the generated proteins have a larger variance in interface area than the original proteins under extensive ablation, which contributes to a relatively high Jensen-Shannon distance between the distributions. However, the divergence shrinks dramatically under moderate and limited ablation. Figure S9 shows that, because uniform ablation removes samples from the entire training distribution evenly, the JS divergence is small on average compared to the corresponding trials in Figure 10. We also note that the generated protein complexes in general do not occupy the same area in latent space as the removed protein complexes (Figure 9).

**Figure 9:**
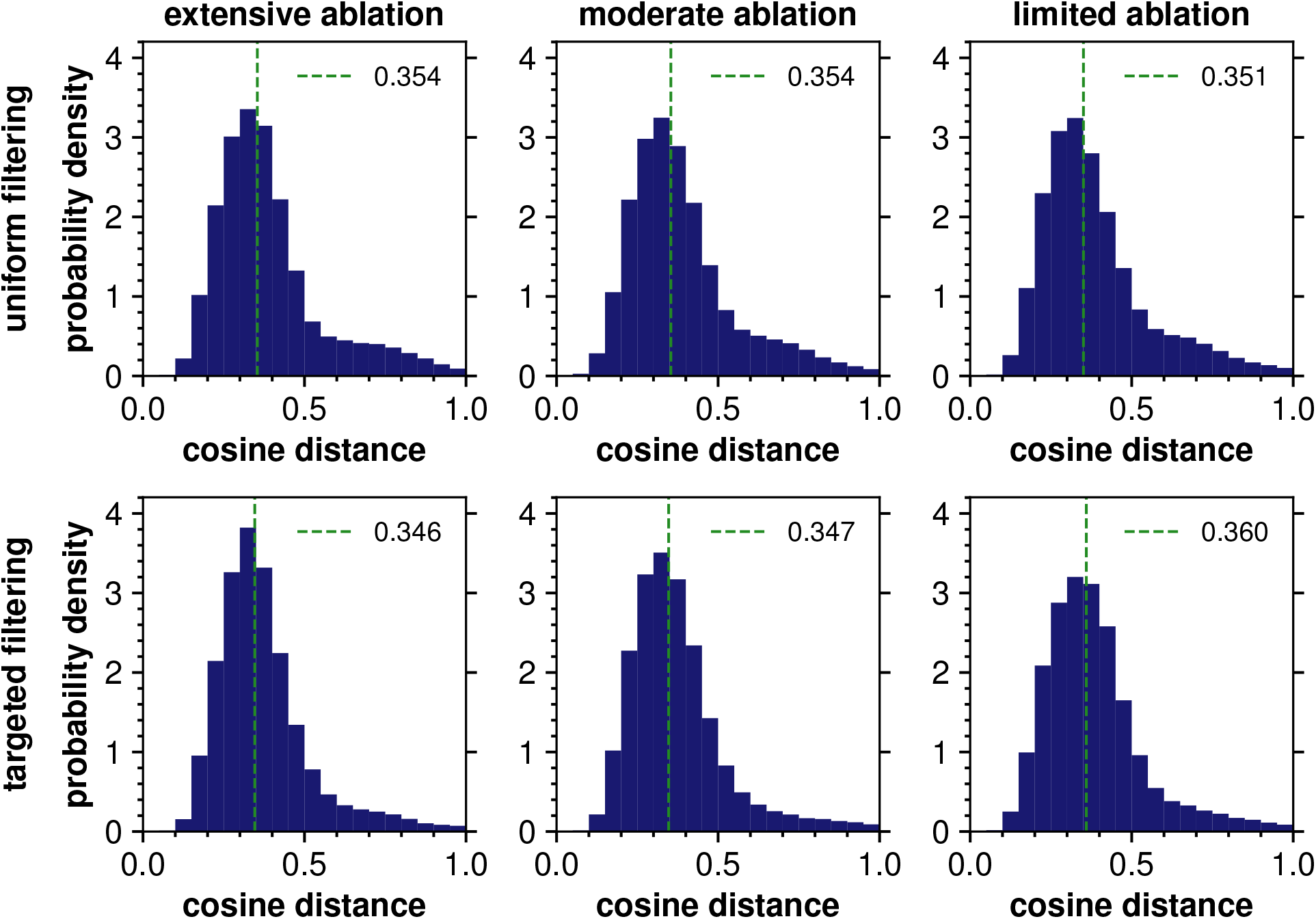
Embedding-space proximity of generated complexes to those removed in ablation. Distributions of cosine distances between generated protein embeddings and ablated protein embeddings for uniform (top) and targeted (bottom) ablation of extensive (left), moderate (middle) and limited (right) degree. Of the five candidates generated for each dissimilar protein seed, the candidate closest to the dissimilar protein embedding is retained for this figure. The median cosine distance is shown by a vertical green line in each panel. Binwidth is 0.05.

**Figure 10:**
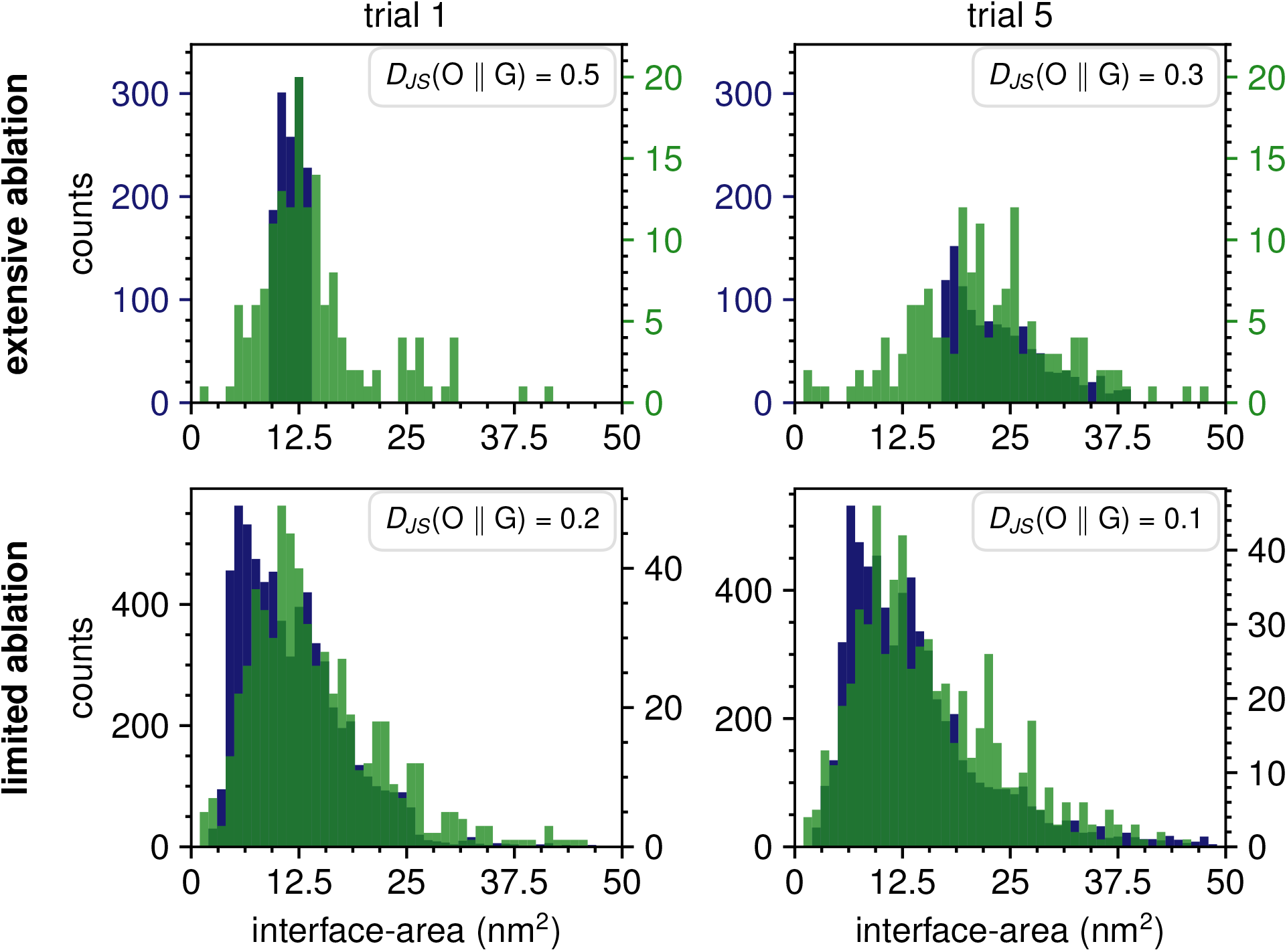
Comparison of interface areas between generated complexes and complexes that remained after ablation. Histograms of the interface areas for generated protein complexes using extensive (top) and limited (bottom) targeted ablation (green) overlaid on the distributions used during initial training (blue). Trials 1 (left) and 5 (right) are shown as representatives for all trials. The Jensen-Shannon distance between the DIPS-Plus proteins and the generated proteins is shown in the top-right corner of each plot. Binwidth is 1 nm^2^.

### Changes in Model Performance When Ablated Sets are Augmented With Generated Samples

We sought to learn whether the complexes we generated could augment the original ablated datasets. Due to the stochastic nature of the protein generation process, we generated five different candidates for each seed (scrambled dissimilar) protein complex and then selected a final candidate for further training, which permitted us to generate on the order of 10^2^ new complexes for each of the ablation conditions. Initial tests comparing training on unaugmented and augmented sets showed no change in train, validation, or test MCC. We note that the ablated sets explored in this work are of sizes on the order 10^3^ *−* 10^4^, so it was possible this lack of change was due to the augmentation not being large enough relative to the un-augmented set.

In light of this, we further reduced the size of the ablated sets by taking a random 10% subsample after the ablation condition was applied. For clarity, we refer to this hereon as “sub-sampling” to not confuse it with the original ablations. For example, the 20% ablations, which normally have 8000 elements, were sub-sampled down to 800, i.e. 2% of the full set. The targeted 20% ablation still only holds samples from within a 20-percentile range of the original DIPS-Plus set, and since the sub-sampling was done uniformly the remaining 800 are uniformly spread throughout that limited range. By that same token the uniform 20% ablation, sub-sampled down to 800 elements, remains spread throughout the full range of interface areas in DIPS-Plus. Then the model using ESM-2 for its twin encoder pair was trained on this set of 800 elements, and trained separately on these 800 augmented with 166 generated proteins and the MCCs compared. Similar statements hold for the other ablations. The results of this approach are shown for targeted and uniform ablations on the interface area variable in Figure 11. The scale of MCC values seen here is substantially smaller than in the prior results on the true 20/50/80% ablations, likely due to the small fraction of training data used. We also see a different trend among ablations that are of different sizes, but belong to the same group of either targeted or uniform ablation. The targeted ablations now clearly improve in performance with size/scope, whereas in prior sections we saw a weak trend of decreasing performance. The uniform ablations maintain their behavior of improving as dataset size increases, even with the 10% sub-sampling.

**Figure 11:**
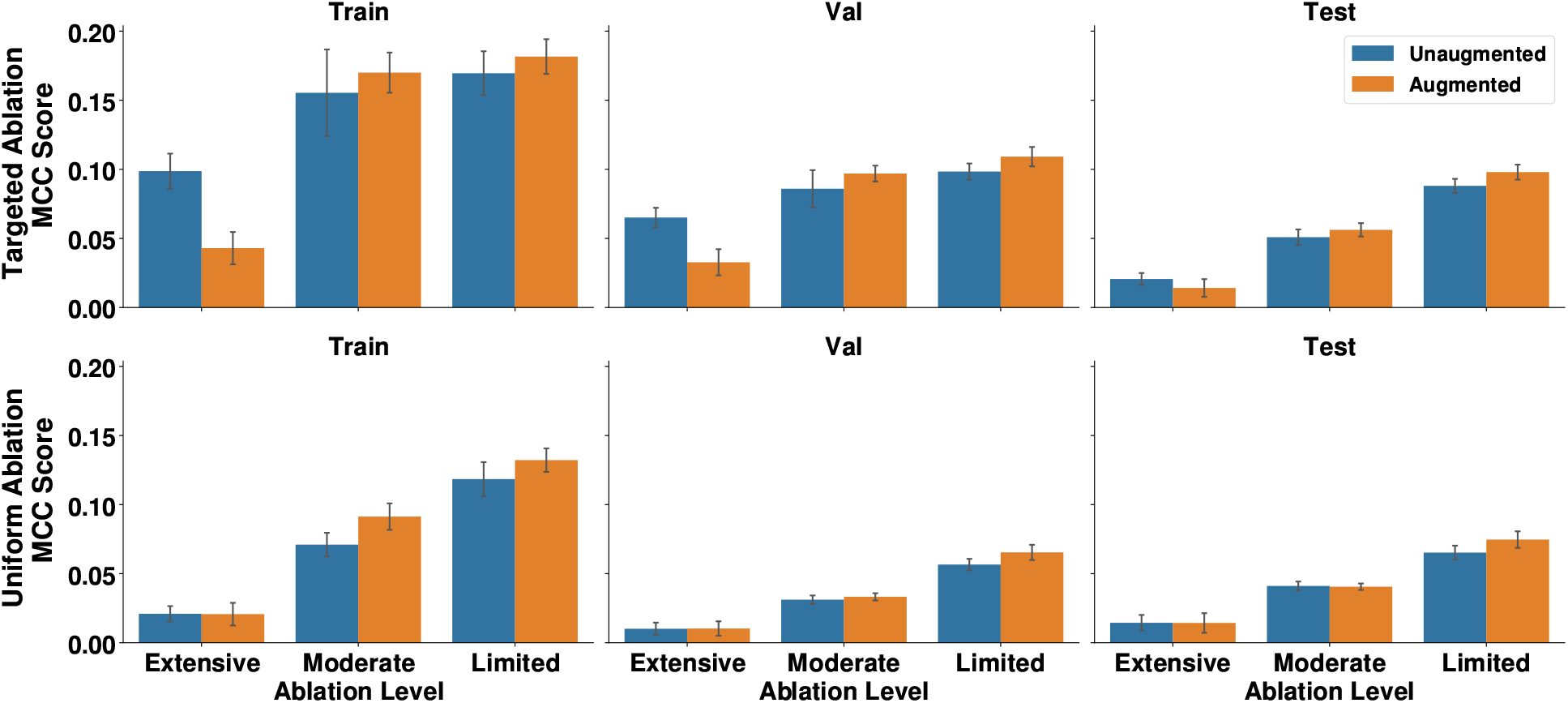
Generative Augmentation Results on ESM-2 Performance for Ablation by Interface Area. Train, Val, and Test MCC scores for ESM-2 on un-augmented (blue) and augmented (orange) targeted (top) and uniform (bottom) ablated sets, with ablation performed using interface area.

Targeted ablations respond better to augmentation, with both 50% and 80% seeing an increase in training, validation, and test MCCs. Uniform ablations, meanwhile, only see improvement upon augmentation when 80% of the set is maintained. Ablated sets of an extremely small size do not respond well to the active learning protocol. 20% ablations fail to improve upon augmentation, with the targeted 20% ablation even seeing a decrease in performance. We note that the corresponding train MCC scores for both the original and the augmented sets are very low, indicative of random predictions.

These trends differ for the sets ablated by residue-residue pair count, shown in Figure S11. In several of these cases the addition of the generated complexes lead to a decrease in performance. Among targeted ablations, only the one that maintained 80% of the dataset showed an improvement in validation or test MCC. This may imply that residue-residue pair count is not as good a metric for focusing the dataset’s distribution as interface area is. We hypothesize that because we used the solvent-accessible surface area (SASA) input track of ESM-3 during the generation process, our protocol is biased in favor of the interface area ablation scheme. There are multiple ways to generate a protein-protein complex with the same interface SASA, but perhaps fewer ways to generate a protein-protein complex with the same interface residue-residue count. The masking process for the dissimilar proteinprotein complexes can change the total number of residues in the protein, which may make generated proteins under the residue-residue scheme less useful for training than generated proteins with a different interface SASA value under interface area ablation.

## Conclusions

### Predictive performance is limited by training data distribution and model size

One of the goals of this project was to investigate how the negative effects of a limited training set size might be mitigated by correspondingly narrowing its distribution. The first step was to establish an upper bound on the performance of a pLM when sufficient data is available. For some sizes of pLMs, DIPS-Plus can be used to locate this upper bound, because DIPS-Plus is a large curated dataset that covers the currently-known biological space of protein complexes. The performance of small models on DIPS-Plus can represent a scenario where one is able to accumulate sufficient data for the needs of a pLM. ESM-2 and AminoBERT were both able to achieve a validation/test MCCs of *∼* 0.5 when trained using the full DIPS-Plus set, unlike ProtBERT which only reached MCCs of *∼* 0.3. ProtBERT was also the largest model used, with 420M parameters compared to only 8M and 87M parameters in ESM-2 and AminoBERT. This is a reminder of the data-hungry nature of pLMs, even among relatively small models that don’t exceed 1B parameters. This problem was only exacerbated when ablating the set to smaller sizes, meaning the results of ProtBERT on these sets were not particularly informative, and it was not incorporated into the AL pipeline discussed in the following section.

Both ESM-2 and AminoBERT demonstrated similar behavior across the ablation trials. The three sets of uniform ablation trials showed MCCs in line with what one might expect for a reduction in dataset size: as the size decreased, MCC across training, validation, and test sets decreased in concert. Validation and test set MCCs were approximately equal to one another across trials, despite hyperparameters having been optimized for validation set performance specifically.

The trials where ablation was performed in a purposefully biased manner showed clear differences in model performance, even though the amount of data seen was the same as in the uniformly ablated trials. The training and validation MCCs were found to be on par with, if not surpassing, those when the full DIPS-Plus set was used. Only the test set MCC showed a substantial decrease in performance relative to both the full set and the uniformly ablated trials. This behavior, coupled with the increase in the train and validation MCCs, is indicative of overfitting. The test MCC also showed a clear trend increasing with respect to the size of the ablated set, while the trends for train and validation MCC are weaker and model-dependent. For instance, ESM-2 saw a weak decrease in train and validation MCCs as the ablated set size increased, while AminoBERT did not.

An additional point that is not reflected in the results plots is that AminoBERT occasionally failed to train on the uniformly ablated sets. These trials trained successfully when reinitialized with a different set of parameters. The smaller ESM-2 model trained successfully on all ablated sets on the first try.

The results on the intentionally biased ablated sets are informative regarding the relationship between a dataset’s size and scope, and their combined effects on how a model learns from it. A training set that is small in size can still be useful to a model, but the amount of information that can be gleaned from it is reduced. Attempting to maintain a distribution akin to the full DIPS-Plus set in a low-data scenario is not ideal, as that information must be spread thinly throughout. If we allow the ablated set to focus its informativeness onto a smaller problem scope, then it still proves to be useful for a model’s learning process.

This usefulness to learning does not extend to inference on an expanded scope, as can be seen by the MCCs on the unablated test set remaining low in the targeted ablations, even when validation MCC is high. In such a case, a uniform distribution remains favorable, even if it still struggles to achieve as high an MCC as was found in the targeted ablation trials.

### Uncertainty-Based Active Learning Requires a Minimum Dataset Size and Performs More Favorably on a Targeted Dataset

The limited ablation test set MCC scores for models trained under interface area ablation increase upon augmentation with the generated protein complexes. However, in every case (uniform vs targeted filtering, interface area vs residue-residue count ablation) the extensive ablation test MCC scores either remain constant or decrease upon augmentation. Additionally, we observe a decrease in MCC scores upon augmentation for both the training and validation sets after interface area and residue-residue count ablation in the extensive ablation scenario. We infer that a lower limit exists, for a given complexity of data and type of model, beneath which augmentation with synthetic data is actually detrimental to improving model performance. If a set of models is given too little training data, that set’s estimation of the uncertainty of data points is unreliable, and the uncertainties are misleading when used to suggest new dataset elements. There may be an upper limit beyond which augmentation is also detrimental for improving model performance. The limited targeted residue-residue count ablation dataset shows a decrease in test set MCC upon augmentation. However, this same decrease is not seen for the limited uniform residue-residue count ablation nor either limited interface area ablations.

Although DIPS-Plus is not a comprehensive dataset of every protein complex, it is the highest quality dataset of such structures currently available. In this study, we already know what the full DIPS-Plus dataset looks like and we know how many proteins we remove from it for each ablation. However, our results suggest a useful rule of thumb for a realistic scenario where the initial amount of data is limited and a “complete” dataset may not be possible to characterize or obtain. In that case, the best strategy for training a predictive ML model is to begin with a targeted core of well-understood data points and then branch out in the space. A targeted strategy restricts the space to which the model applies, which makes identifying additional data points to gather or generate a more reliable process. This approach minimizes the out-of-distribution error associated with a thinly-spread set of data across a whole measure space, which implies that any local region of that space is likely poorly understood. Other work has demonstrated similar results. Endo and Kaneko explored Bayesian optimization (BO) in small-data scenarios applied to reaction condition optimization.^37^ BO is closely related to AL, attempting to find the optimum of an unknown function by iteratively sampling points from its domain.^38^ They made the argument that one finds greater success in such scenarios by optimizing not exact prediction of reaction yield of particular conditions, but instead the simpler task of ranking the conditions from best to worst.^37^

Our results on the interface area distribution of the complexes we generated using ESM-3 indicated a bias of that model towards areas in the range of 0-25 nm^2^, where the majority of wild-type protein-protein complexes are found. This is possibly due to the nature of its finetuning, where it was trained to reconstruct existing proteins and treat those as characteristic of any protein. Hou et al. studied the performance of similarly finetuned pLMs in mutation fitness prediction, and found that models could become overly fixed in its conception of what is possible for a protein, being more likely to score any deviation from its training set as deleterious.^39^ We hypothesize this may be occurring in our generative model, implying that such models may be restricted in their tendency to produce novel proteins that fall beyond what they have already seen in their pre-training. The creators of ESM-3 explored the capacity of their model to generate proteins distinct from their pretraining data, and found that doing so requires deliberately crafted prompting.^16^

### Limitations and Future Directions

We were limited in this work to only using one dataset and three pLMs, with only two of those able to finetune well enough for use in the downstream AL protocol. DIPS-Plus, while large and diverse, does not cover the full diversity of protein-protein complexes, focusing primarily on those stable enough to have their atomic structure well-resolved. There also exist AL protocols more sophisticated than what was presented here and whose performance in low-data scenarios with pLMs we are unable to draw conclusions on.

As the capacity of available models continues to grow, the limitations of data collection for chemical domains is liable to become more and more problematic. Additional studies on other datasets, problem types, and AL protocols will be helpful to determine the best procedures for modern machine learning in the chemical sciences. Generative models are a promising avenue for data augmentation without requiring bench work, but have their own in-built biases due to the nature of their training that may require increased care for successful use.

## Acknowledgement

Research reported in this publication was supported by the National Science Foundation grants MCB-2527485 (to R.I.D) and MCB-2136816 (to G.S.), and by the National Institute of Biomedical Imaging and Bioengineering of the National Institutes of Health under grant number R01EB035594 (to P.S.). This work used the ACCESS program’s advanced computational resources through the allocation BIO210094 (to R.I.D.) and allocation MCB170020 (to G.S.).

The content is solely the responsibility of the authors and does not necessarily represent the official views of the National Institutes of Health.

## Data and Software Availability

The code needed to reproduce the results in this paper is available on GitHub at https://github.com/DimaUClab/protein-complex-ml. The data collected and discussed in this paper are available on Zenodo at 10.5281/zenodo.23453970. Because the predicted contact maps predicted for each trial total over 1.5 TB of storage, they are available from the authors upon request.

## Supporting Information Available

The Supplementary Information provides the following:

• **T1** Hyperparameters for fine-tuning the AminoBERT, ProtBERT, and ESM-2 models

• **S1** Comparison of contact map similarity scoring functions

• **S2** Average number of masked residues percent of complex masked after masking according to Table 3

• **S3** MCC scores for models trained under all residue-residue count ablations

• **S4** CKA-scores of protein-protein complexes according to models trained under residue-residue count ablation

• **S5** CKA-score skewnesses for models trained under interface area ablations

• **S6** CKA-score skewnesses for models trained under residue-residue count ablations

• **S7** Histograms of interface areas for all targeted ablations with low and high CKAscores

• **S8** Histograms of interface areas for all uniform ablations with low and high CKAscores

• **S9** Representative histograms of interface areas of generated protein complexes under uniform ablation

• **S10** Representative histograms of the cosine distance distributions of similar and dissimilar protein complexes after residue-residue count ablation

• **S11** MCC scores after augmentation with generated protein complexes after residueresidue count ablations

---

**TOC Graphic.**
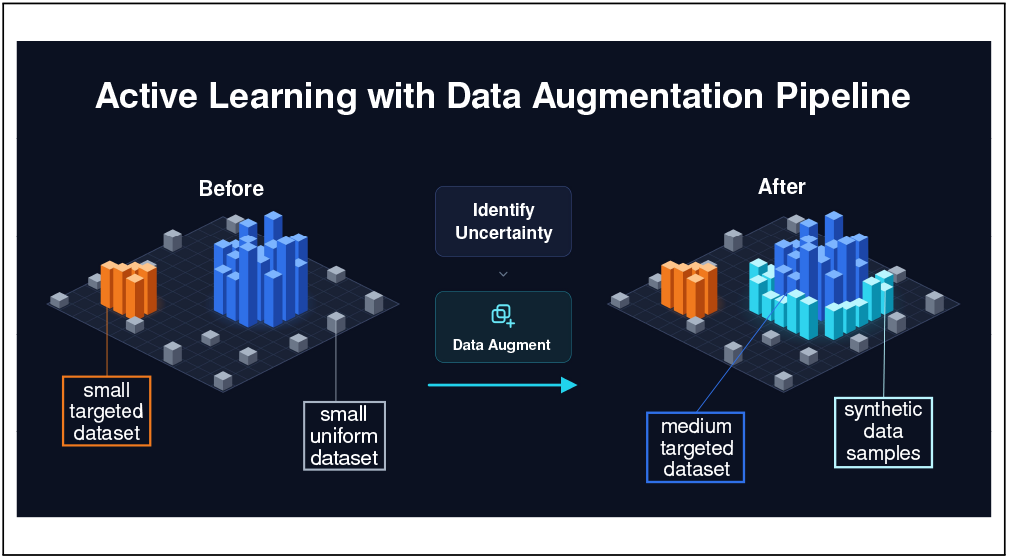

