## supplemental tables and figures for "Building from the ground up: augmentation strategies to improve protein language model performance on protein sets"

[b]co-first authors

[c]co-corresponding authors

Table S1: The variables and their values explored during hyperparameter optimization. Square brackets are used to denote continuous ranges and curly braces are used to denote discrete sets. Log-uniform sampling was used for the learning rate, while all other hyperparameters were sampled uniformly.

| Hyperparameter | Values Explored |
|---|---|
| Batch Size | {4,8} |
| Learning Rate | [1e-5, 1e-3] |
| # Epochs | {5, 10, 15} |
| MLP Depth | {2, 3, 4} |
| MLP Width | {32, 64, 128} |
| MLP Dropout Rate | [0.0. 0.5] |
| MLP Activation Function | {ReLU, GELU, Tanh} |
| BCE Position Weight | [1.0, 100.0] |

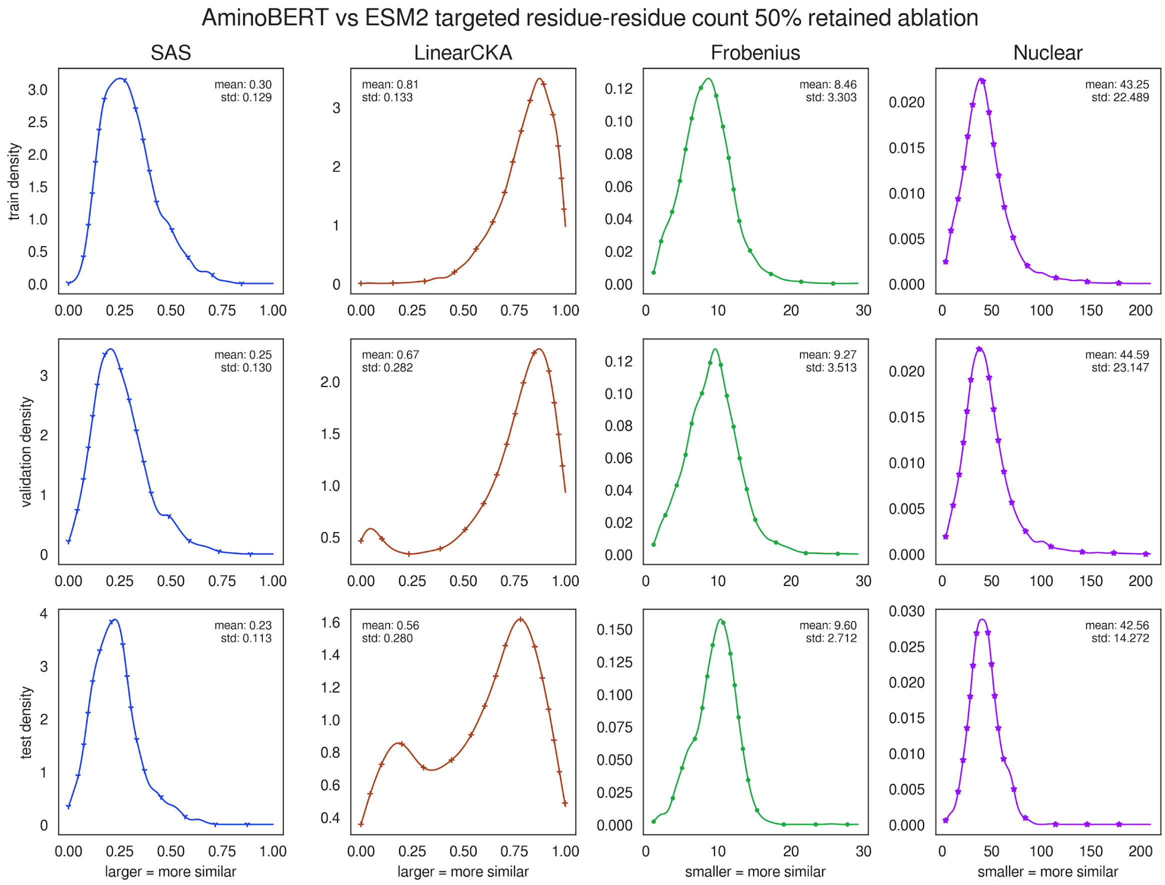


Figure S1. AminoBERT vs ESM-2 similarities for the targeted moderate residue-residue count ablation. For each metric, the distribution of similarities is plotted for the train, validation, and test sets. The (linear) CKA metric produces distributions with a higher standard deviation than Singular Angle Similarity (SAS) and better differentiates the train, validation, and test sets compared to the Frobenius and Nuclear metrics. CKA and SAS similarity increases with increasing value, while the Frobenius and Nuclear similarity decreases with increasing value.

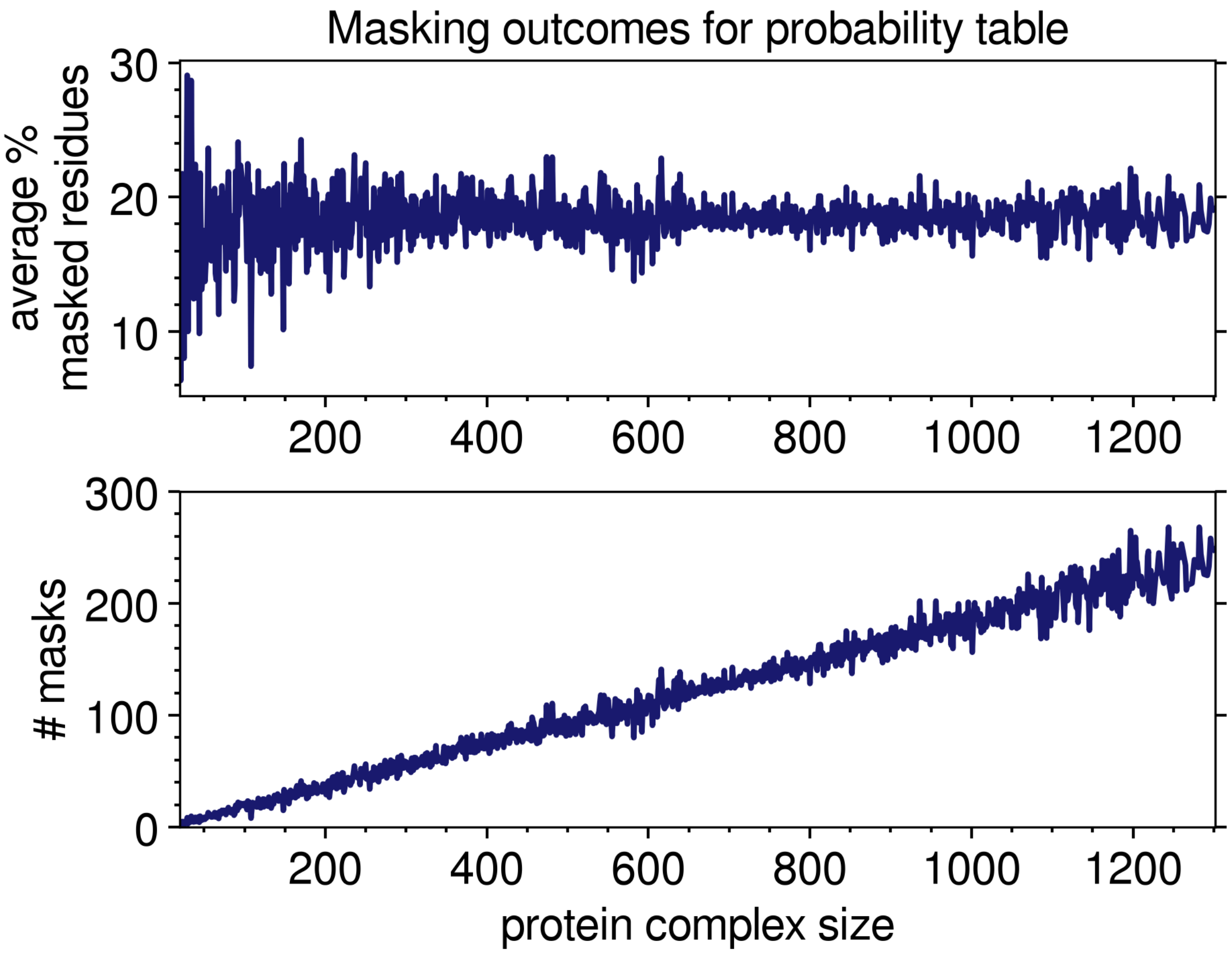


Figure S2. The average percent of residues masked (upper plot) and average number of mask tokens (lower plot) for each protein complex size. On average 20% of the residues for a protein-protein complex are masked, and the average number of masks linearly increases with increasing protein complex length.

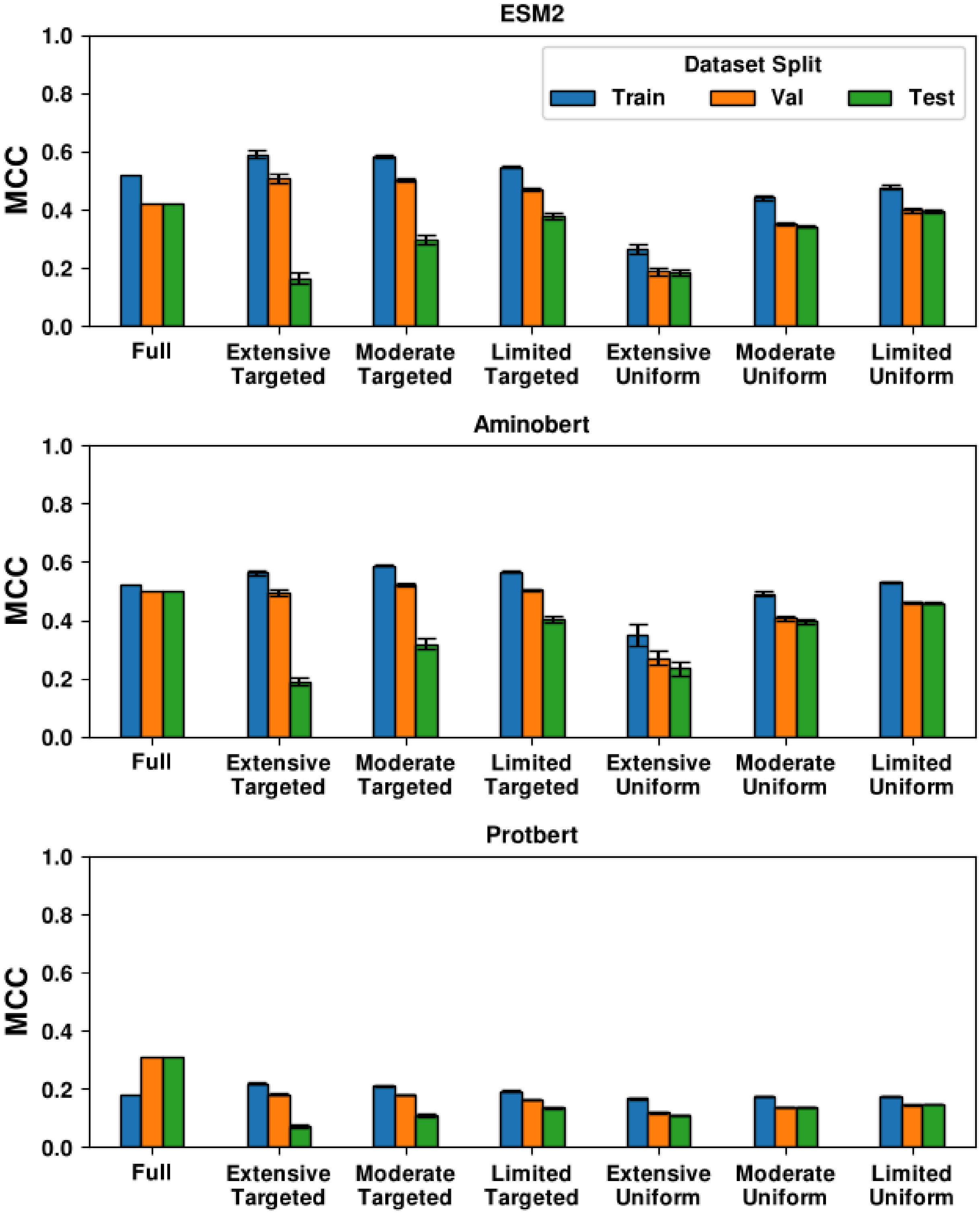


Figure S3. Performance of pLMs on DIPS-Plus data. Train (blue), validation (orange), and test (green) MCCs for ablation trials performed using residue-residue pair count, averaged across the five trials tested for each ablation condition.

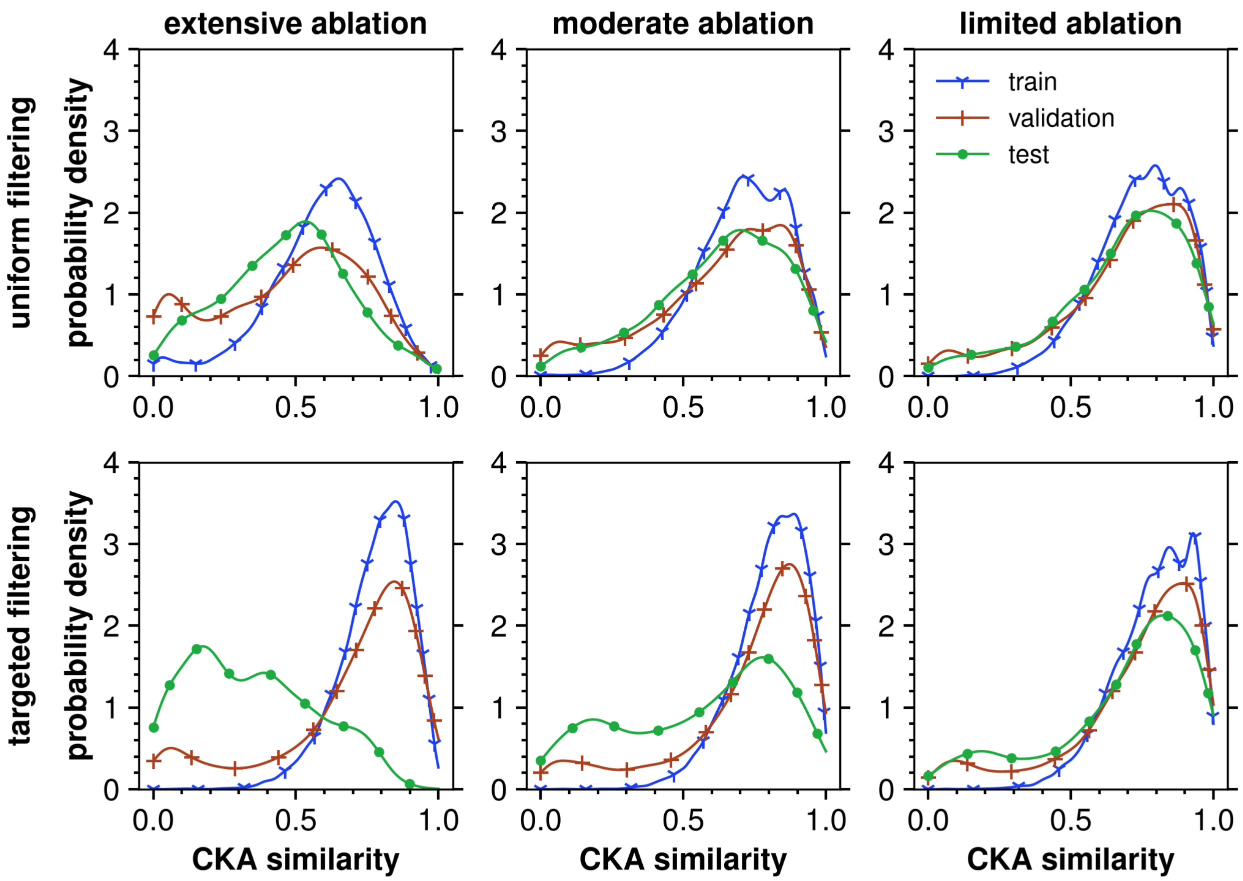


Figure S4. AminoBERT vs ESM-2 residue-residue count ablation similarities. Gaussian kernel density plots of the CKA-scores under each ablation method (uniform or targeted) and ablation degree (extensive, moderate, or limited) after ablation under the residue-residue count criterion. Models trained after uniform ablation (top row) exhibit less agreement on average (smaller CKA scores) than models trained after targeted ablation (bottom row). However, the differences between uniform and targeted ablation are strongest when very little data remains in the dataset (column 1) and almost disappear when most of the dataset remains (column 3).

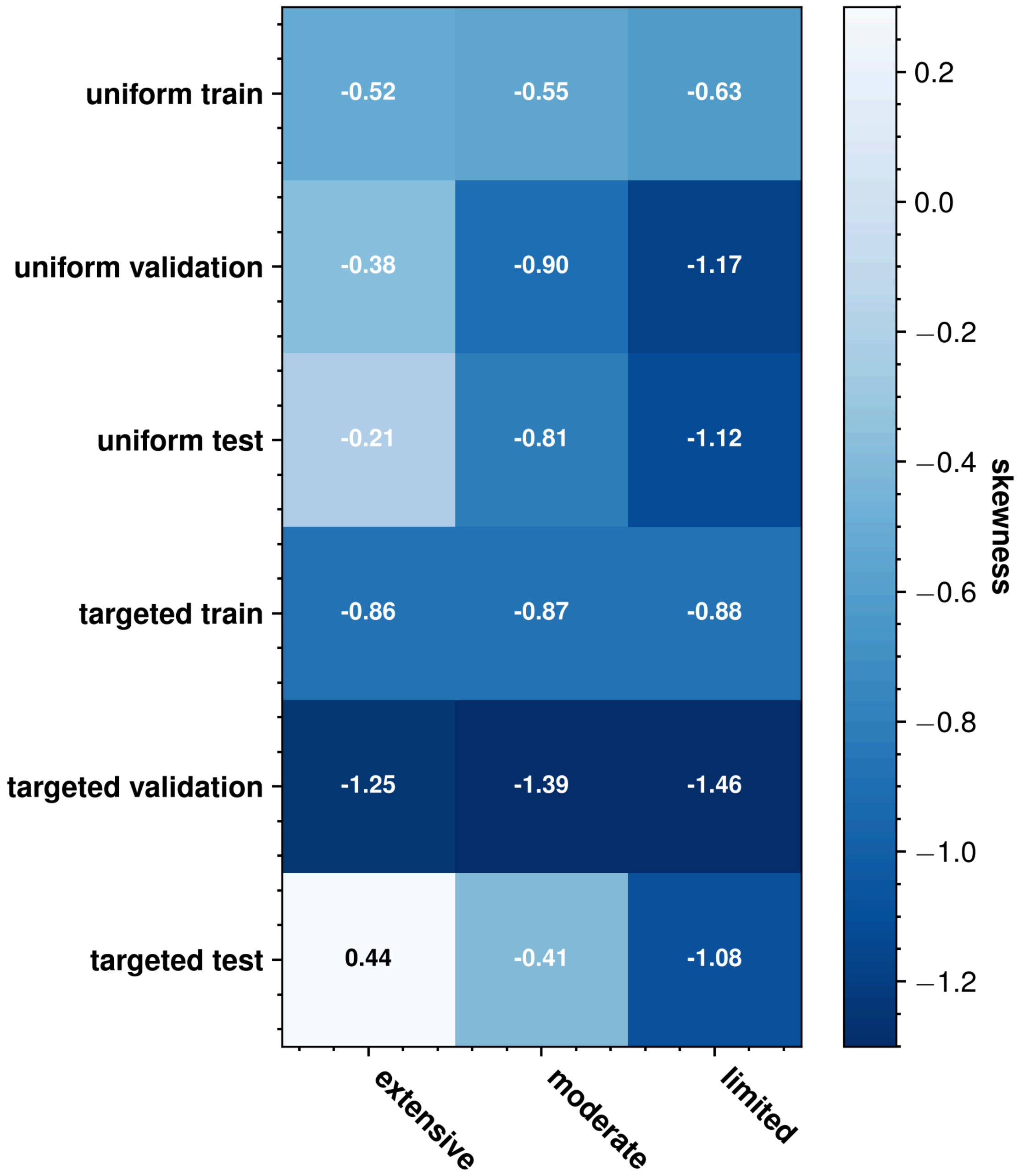


Figure S5. Skewness of the CKA-score distributions for the training, validation, and test sets under interface-area ablation. Negative skewness indicates more of the probability density than a normal distribution sits to the left of the mean, while positive skewness indicates more probability density than a normal distribution sits to the right of the mean. Models trained on the targeted extensively ablated dataset have very little overlap in their contact map predictions on the test set, which is the only distribution with positive skewness.

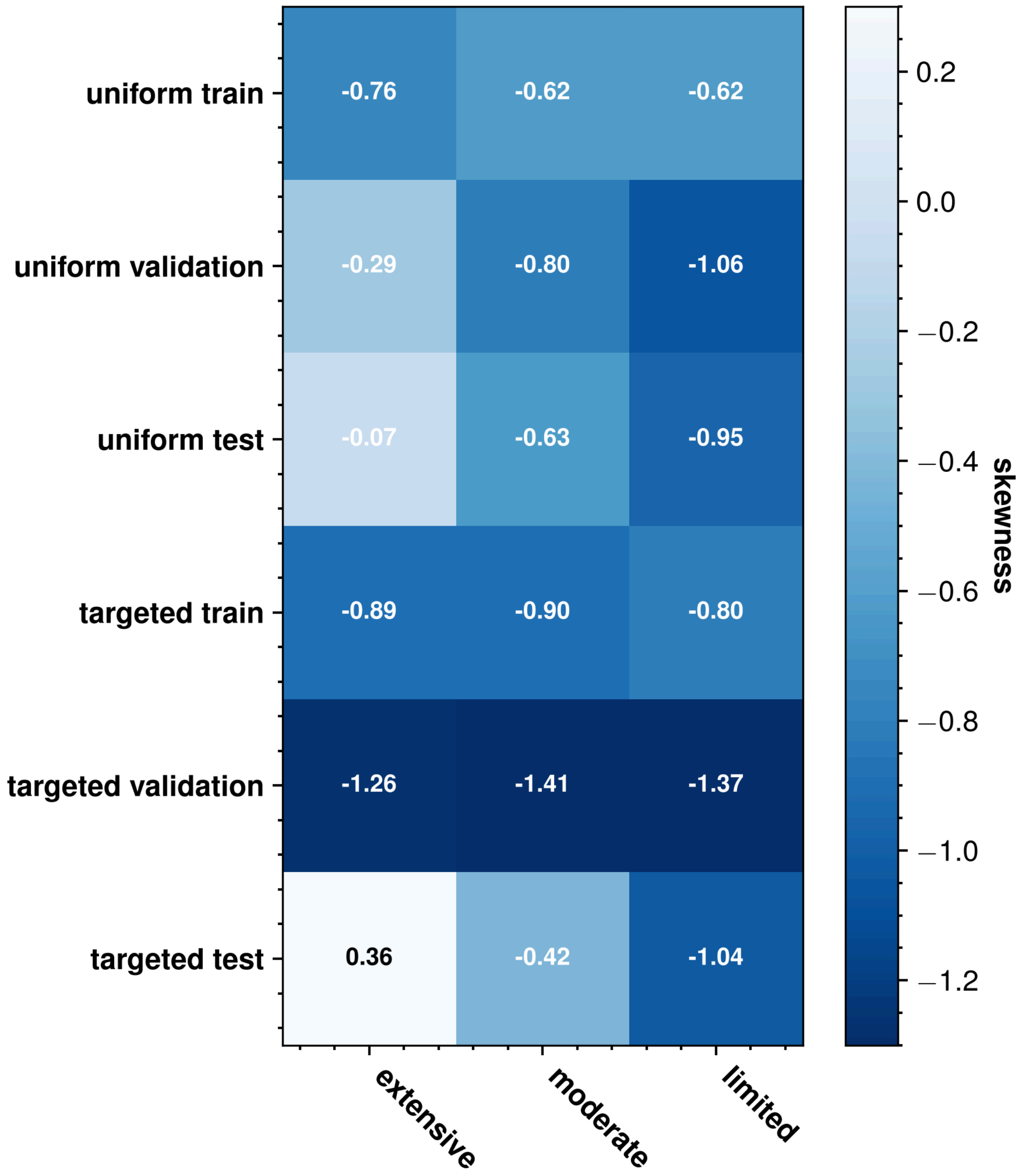


Figure S6. Skewness of the CKA-score distributions for the training, validation, and test sets under residue-residue count ablation. Negative skewness indicates more of the probability density than a normal distribution sits to the left of the mean, while positive skewness indicates more probability density than a normal distribution sits to the right of the mean. Models trained on the targeted extensively ablated train set have very little overlap in their contact map predictions on the test set, which is the only distribution with positive skewness.

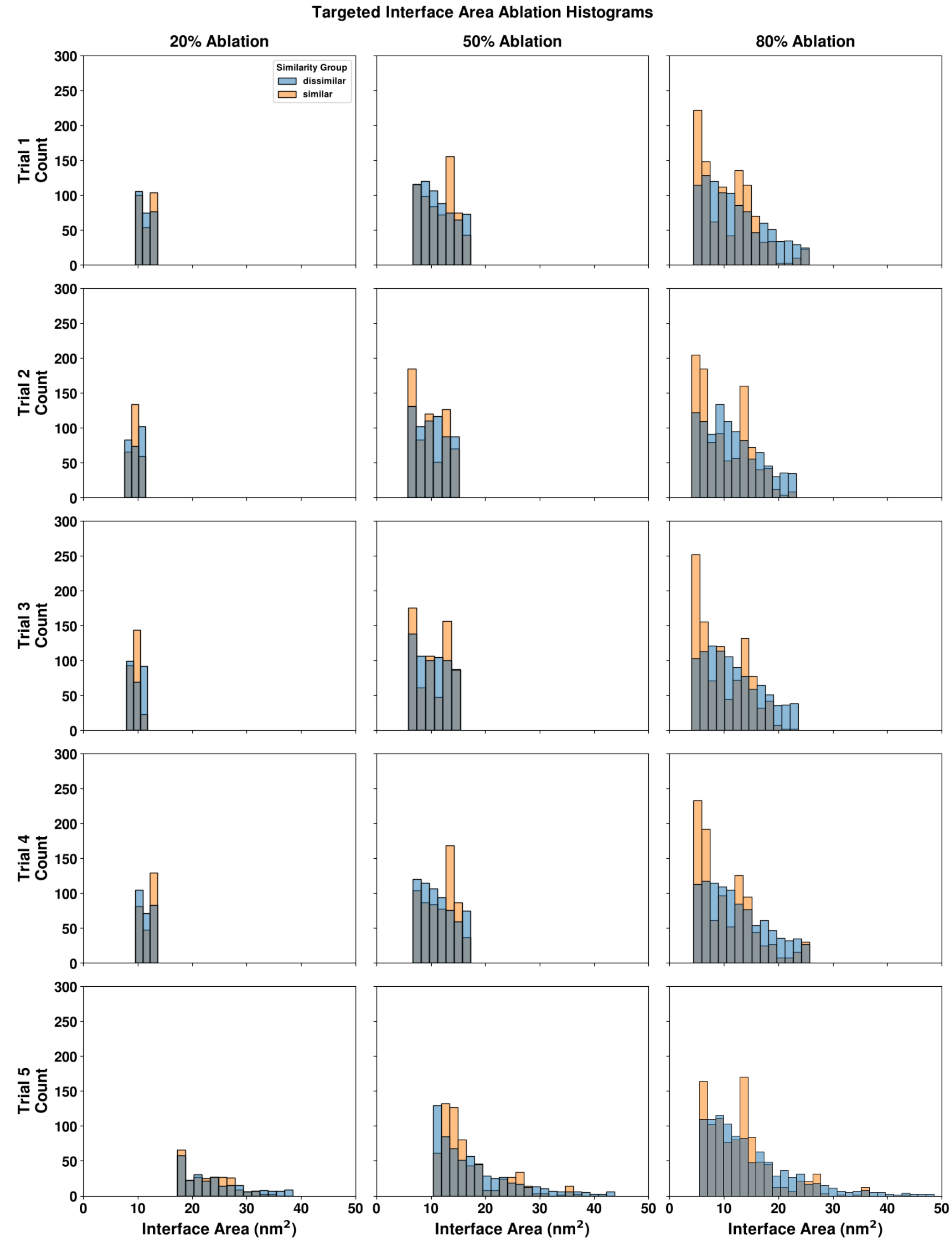


Figure S7. Interface area distributions for all targeted ablated DIPS-Plus sets. Comparison of similar (orange) and dissimilar (blue) interface area distributions for all 30 targeted ablation trials, with five trials (separated by row) done for each of extensive (left), moderate (middle) and limited (right) Binwidth was set to $1nm^2$ for all plots.

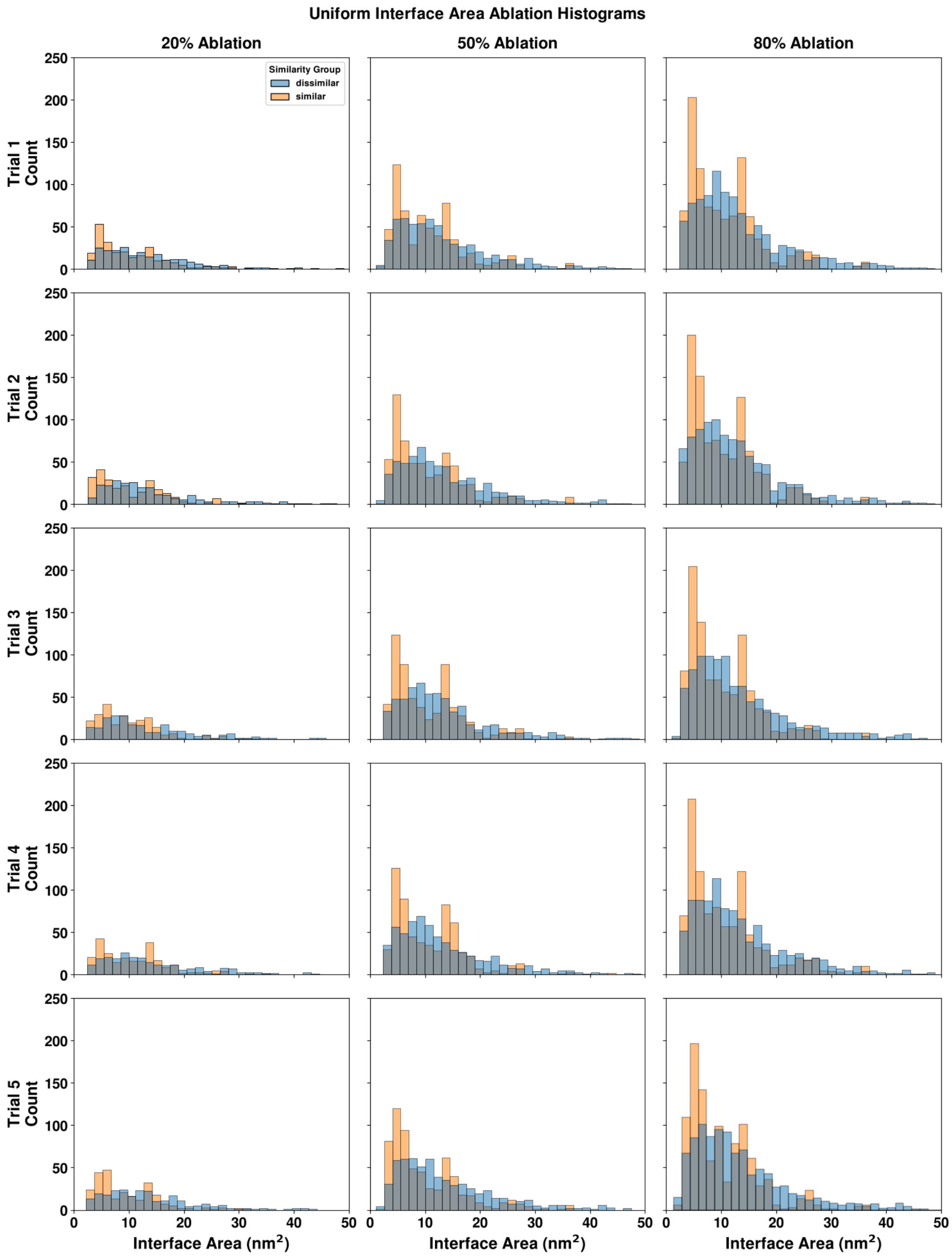


Figure S8. Interface area distributions for all uniform ablated DIPS-Plus sets Comparison of similar (orange) and dissimilar (blue) interface area distributions for all 30 uniform ablation trials, with five trials (separated by row) done for each of extensive (left), moderate (middle) and limited (right) Binwidth was set to 1nm$^2$ for all plots.

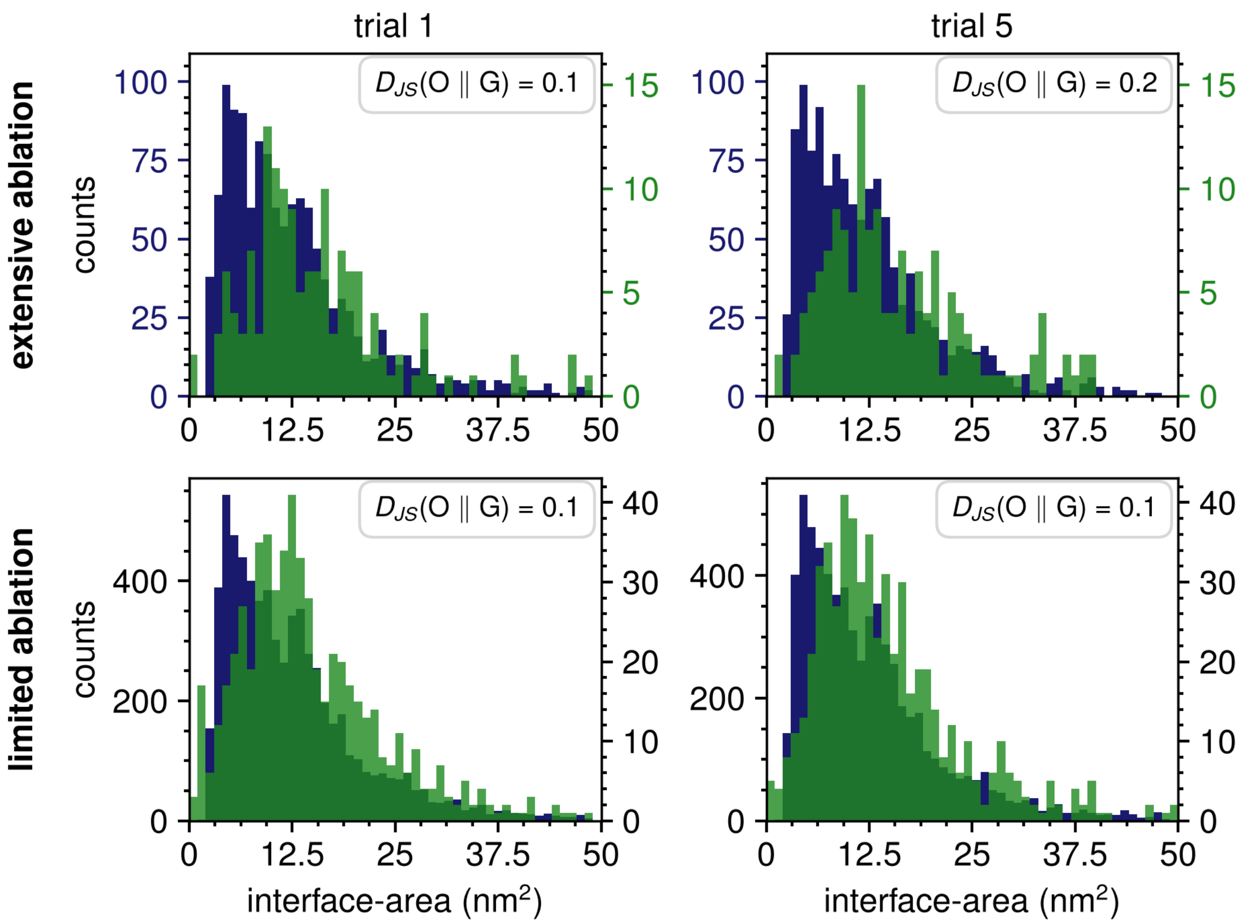


Figure S9. Histograms of the interface-areas for generated protein complexes (green) overlaid on the distributions used during initial training (blue) after uniform ablation. Trials 1 and 5 are shown as representatives for all trials. The Jensen-Shannon (JS) distance between the DIPS-Plus proteins to the generated proteins is shown in the top-right corner of each plot. Binwidth is 1 nm$^2$.

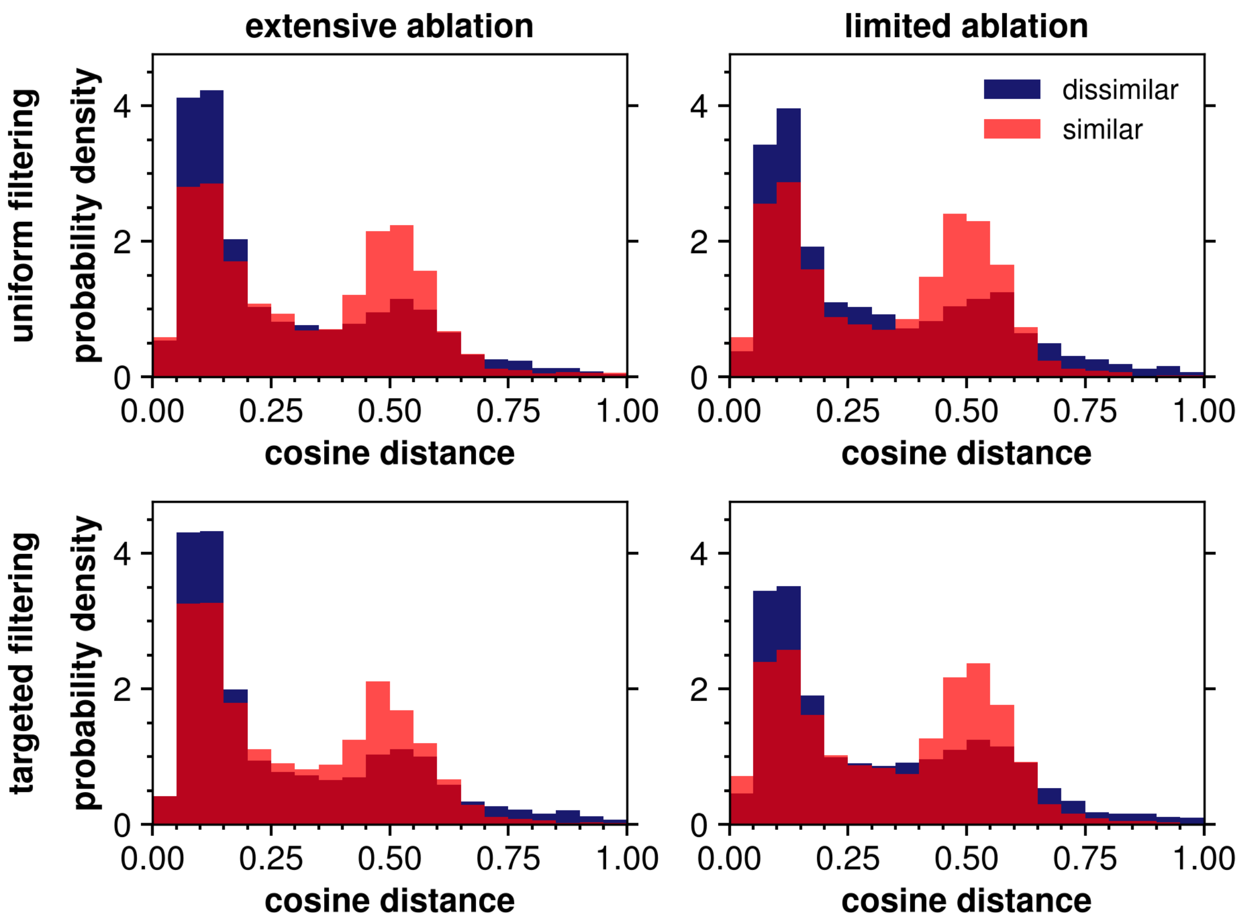


Figure S10. Distributions of cosine distances between dissimilar (blue) and similar (red) protein complexes vs removed proteins under residue-residue count ablation. The dissimilar complexes are closer overall to the ablated complexes than the similar complexes. Binwidth is 0.05. The embeddings were obtained from the ESM-C 6B model.

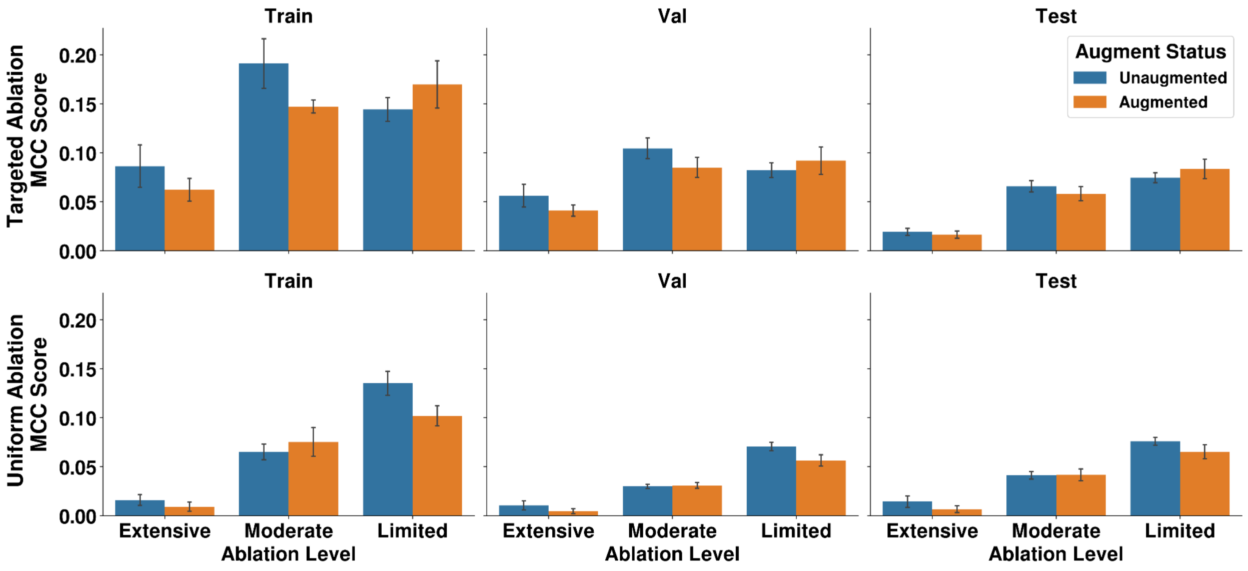


Figure S11. Generative Augmentation Results on Model Performance for Ablation by Residue-Residue Pair Count. Train, Val, and Test MCC scores for ESM-2 on un-augmented (blue) and augmented (orange) on biasedly (top) and uniformly(bottom) ablated sets.
